# StreptoCAD: An open-source software toolbox automating genome engineering workflows in streptomycetes

**DOI:** 10.1101/2024.12.19.629370

**Authors:** Lucas Levassor, Christopher M. Whitford, Søren D. Petersen, Kai Blin, Tilmann Weber, Rasmus J. N. Frandsen

**Author notes:** = to whom correspondence should be addressed: Tilmann Weber, Rasmus J.N. Frandsen.

## Abstract

Streptomycetes hold immense potential for discovering novel bioactive molecules for applications in medicine or sustainable agriculture. However, high-throughput exploration is hampered by the current *Streptomyces* genetic engineering methods that involve the manual design of complex experimental molecular biological engineering strategies for each targeted gene. Here, we introduce <u>StreptoCAD</u>, an open-source software toolbox that automates and streamlines the design of genome engineering strategies in *Streptomyces,* supporting various CRISPR-based and gene overexpression methods. Once initiated, StreptoCAD designs all necessary DNA primers and CRISPR guide sequences, simulates plasmid assemblies (cloning) and the resulting modification of the genomic target(s), and further summarizes the information needed for laboratory implementation and documentation. StreptoCAD currently offers six design workflows, including the construction of overexpression libraries, base-editing, including multiplexed CRISPR-BEST plasmid generation, and genome engineering using CRISPR-Cas9, CRISPR-Cas3, and CRISPRi systems. In addition to automating the design process, StreptoCAD further secures compliance with the FAIR principles, ensuring reproducibility and ease of data management via standardized output files. To experimentally demonstrate the design process and output of StreptoCAD, we designed and constructed a series of gene overexpression strains in *Streptomyces Gö40/10*, underscoring the tool’s efficiency and user-friendliness. This tool simplifies complex genetic engineering tasks and promotes collaboration through standardized workflows and design parameters. StreptoCAD is set to transform genome engineering in *Streptomyces*, making sophisticated genetic manipulations accessible for all and accelerating natural product discovery.

**Graphical abstract:** 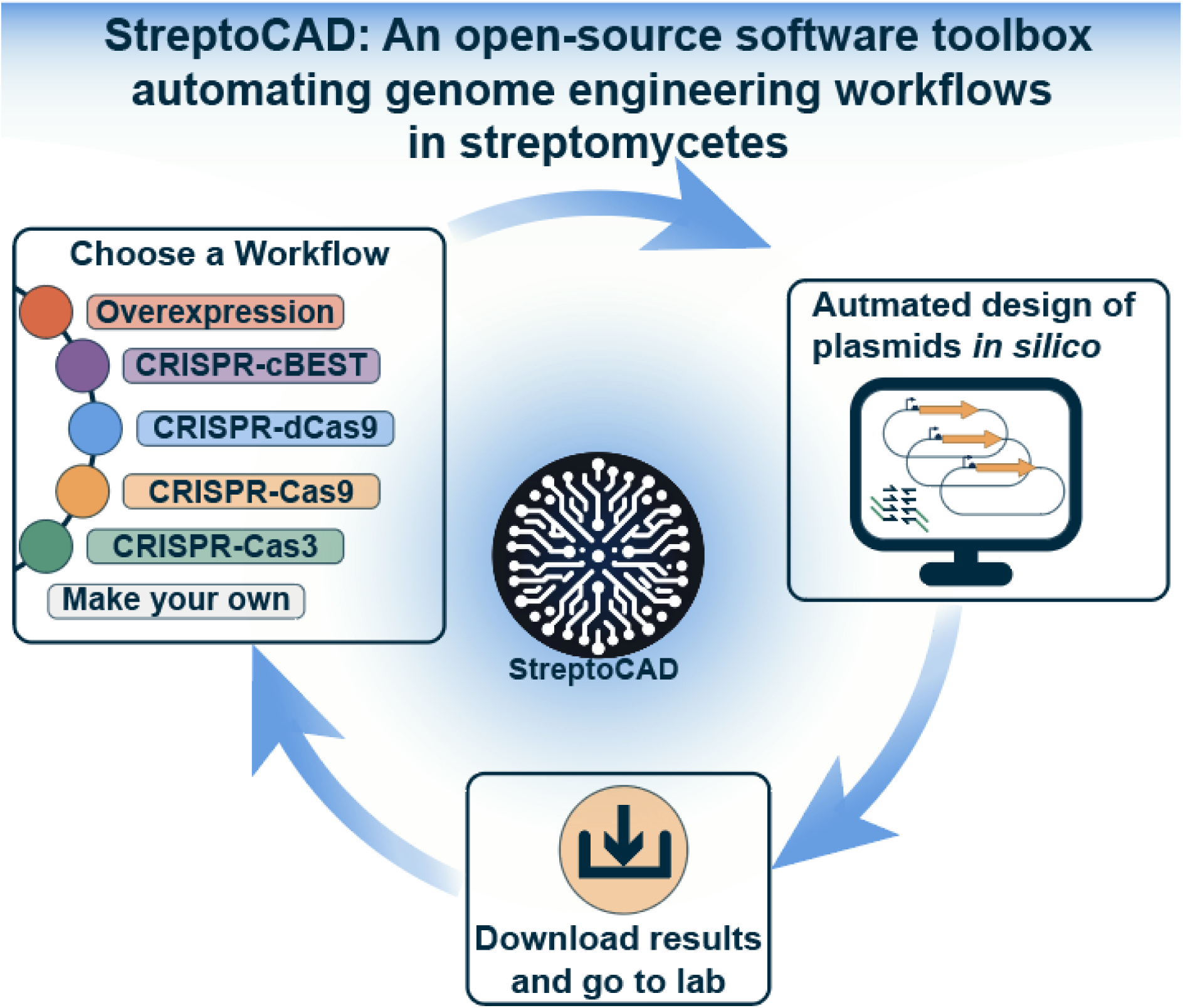

## Introduction

Biotechnology holds the potential to transform the production of foods, materials, fuel, and medicines, offering promising solutions for planetary health and sustainability^1^. A key aspect of this transformation is the discovery and utilization of natural products, which are chemical compounds produced by living organisms with applications in medicine, agriculture, and the chemical industry. In this context, soil and marine-dwelling actinomycetes are of particular interest. These gram-positive bacteria are known to harbor a remarkable number of biosynthetic gene clusters (BGCs) for the production of bioactive specialized metabolites^2,3^. Notably, streptomycetes, the largest genus of actinomycetes, has been a prolific source of new medicines, including antibiotics such as cephamycin, chloramphenicol, and tetracycline; the antifungal cyphomycin; the antiviral compound virantmycin B; immunomodulators like FK506; and anti-cancer medications such as doxorubicin and bleomycin, which have had significant impacts on treating various diseases^4,5^. Beyond therapeutics, streptomycetes have also been a source of agrochemicals such as pteridic acids, which can alleviate abiotic stress in plants^6^ thereby increasing the harvest yields.

The potential of *Streptomyces* for natural product discovery remains vast, offering a promising avenue for developing novel therapeutic and agricultural agents. While genome mining has revealed a vast untapped potential, engineering streptomycetes remains challenging, limiting efficient investigation of the unexplored bio-chemical space^7^. Establishing gene/BGC-function relationships typically involves constructing numerous genetically engineered strains to test various hypotheses^8^. To enhance our understanding, it is necessary to develop new methods that allow for the rapid and efficient construction of strains. Insights from other well-studied organisms, such as *Saccharomyces cerevisiae* and *Escherichia coli*, can guide this effort. For example, Hossain *et al.* 2020 characterized a total of 4350 nonrepetitive bacterial promoters and 1722 non-repetitive yeast promoters with varying transcription rates^9^. This level of characterization and strain construction surpasses recent studies on *Streptomyces* by more than an order of magnitude^10–12^. Moreover, the filamentous growth and complex life cycle of *Streptomyces* further complicate larger-scale automation, presenting design and throughput challenges due to their morphology and lifestyle.

Advancements in genome editing and engineering techniques for streptomycetes, including CRISPR-Cas9, CRISPR base editing (e.g., CRISPR-BEST), CRISPRi, CASCADE-Cas3, and CRISPRa, have increased the efficiency of genetic modifications and cut down the time of building strains significantly^13–18^. However, applying these techniques on a high-throughput scale poses significant challenges^19^. Currently, much of molecular biology work, such as primer design, is done manually. This manual process is error-prone and time-consuming, especially when scaled to handle tens, hundreds, or thousands of strain designs due to the intricate steps involved. As the complexity of designs increases, computer-aided design (CAD) software becomes crucial for minimizing errors, accelerating construction, and standardizing design parameters. By automating tasks such as primer and plasmid design, and simulating assemblies, CAD software enhances experimental consistency and efficiency. This automation facilitates rapid hypothesis testing and enables the generation of large, high-quality datasets for improved learning and research outcomes.

CAD software tools such as Snapgene^20^, CLC-Workbench^21^, and Teselagen^22^ excel in assembling plasmids and simulating parts of experiments within the Design-Build-Test-Learn (DBTL) cycle. However, these tools often lack the flexibility needed to integrate new methods or workflows developed in fast-paced research environments and automate workflows. Additionally, while Teselagen offers some degree of automation and workflow capabilities, tools like SnapGene provide limited options for automating design processes, which can restrict scalability and efficiency. Their "click-and-drop" interfaces, while user-friendly, offer limited standardization, traceability, and throughput. In contrast, open-source tools are typically more adaptable and can be customized to meet specific research needs. Python, in particular, provides a highly flexible platform for developing these tools, with extensive libraries and frameworks that facilitate rapid development and integration of new functionalities. However, these open-source tools often come with a steeper learning curve, which may pose challenges for users without extensive programming experience. Examples of open-source software tools for executing full workflows include GalaxySynbioCAD^23^ and Galaxy^24^. Briefly, GalaxySynbioCAD builds on the Galaxy platform, providing tools specifically tailored for synthetic biology, including retrosynthesis and metabolic pathway analyses throughout the DBTL cycle. The Galaxy platform offers extensive bioinformatics workflows for genomic analysis through its user-friendly web interface.

Despite the customization and flexibility these CAD tools offer, a gap persists in tools that fully address the unique requirements of *Streptomyces* engineering. Researchers often rely on multiple specialized tools to meet their needs, as no single tool currently offers a comprehensive end-to-end solution. For instance, tools like antiSMASH^25^, DeepBGC^26^, and others are widely used to detect biosynthetic gene clusters in *Streptomyces*. However, to design genetic engineering experiments to experimentally analyze the identified BGCs, the research needs to utilize other software solutions such as CRISPy-web^27,28^, CRISPOR^29^, and TransCRISPR^30^ that allow for designing single guide RNAs (sgRNAs). Once the sgRNAs are identified, a CAD software platform like Benchling^31^, or the previously mentioned CAD software tools, can be used to assemble the plasmid targeting the desired genomic region. The use of multiple task-specific tools in *Streptomyces* engineering reflects the fragmented nature of current design workflows, with each tool addressing a distinct aspect of the process. Transitioning bioengineering from an artisanal to a standardized practice requires embracing FAIR (Findable, Accessible, Interoperable, and Reusable) principles. Achieving this will necessitate developing new tools and platforms that seamlessly integrate with existing technologies while providing the flexibility to accommodate evolving research needs and workflows.

In this work, we introduce StreptoCAD, a software toolbox designed to support genome engineering workflows in *Streptomyces*. StreptoCAD automates key aspects of the genome engineering process, by automating the entire design process and the simulation of correct assemblies in *Streptomyces*. StreptoCAD thereby eliminates repetitive manual tasks, speeds up the design steps of the engineering cycle, and facilitates the generation of more strains following a standardized design paradigm for consistent and more reproducible designs, which combined improves the potential for learning and in the future may provide an interface to automation^32^. We experimentally validated this workflow through the successful overexpression of transcriptional regulators in *Streptomyces* sp. Gö40/10, without DNA-level redesigns or experimental errors. Furthermore, by standardizing these workflows, StreptoCAD enhances the ability to share and troubleshoot lab protocols more efficiently, promoting collaboration and consistency in the field.

## Results

### Streamlining genome engineering in *Streptomyces* with StreptoCAD

StreptoCAD is designed to simplify genome engineering by providing out-of-the-box software solutions that streamline tasks such as primer and plasmid design (**Figure 1A**). By building on established protocols and molecular biology techniques, StreptoCAD minimizes the often time-consuming and error-prone nature of manual steps in the design processes, such as copying, pasting, or highlighting sequences across different software applications. The StreptoCAD hence supports more efficient use of time and resources, facilitates faster research advancements, and ensures that all data is saved for later retrieval and analysis, aligning with FAIR data management principles. StreptoCAD offers six distinct workflows for constructing various types of genetically engineered strains (**Figure 1A**), each accompanied by a video illustrating its use (**Suppl. Figure 1-6**). We first developed StreptoCAD Workflow 1, a novel overexpression workflow, and experimentally validated the functionality of the designed materials (oligonucleotides) and predictions (resulting plasmids and strains). Building upon the code elements of Workflow 1, we subsequently developed StreptoCAD Workflows 2–5, which are based on the CRISPR-Cas protocols developed by Tong *et al.* 2015^33^, Tong *et al.* 2019^13^, Tong *et al.* 2020^15^, and Whitford *et al.* 2023^18^ respectively.

**Figure 1:**
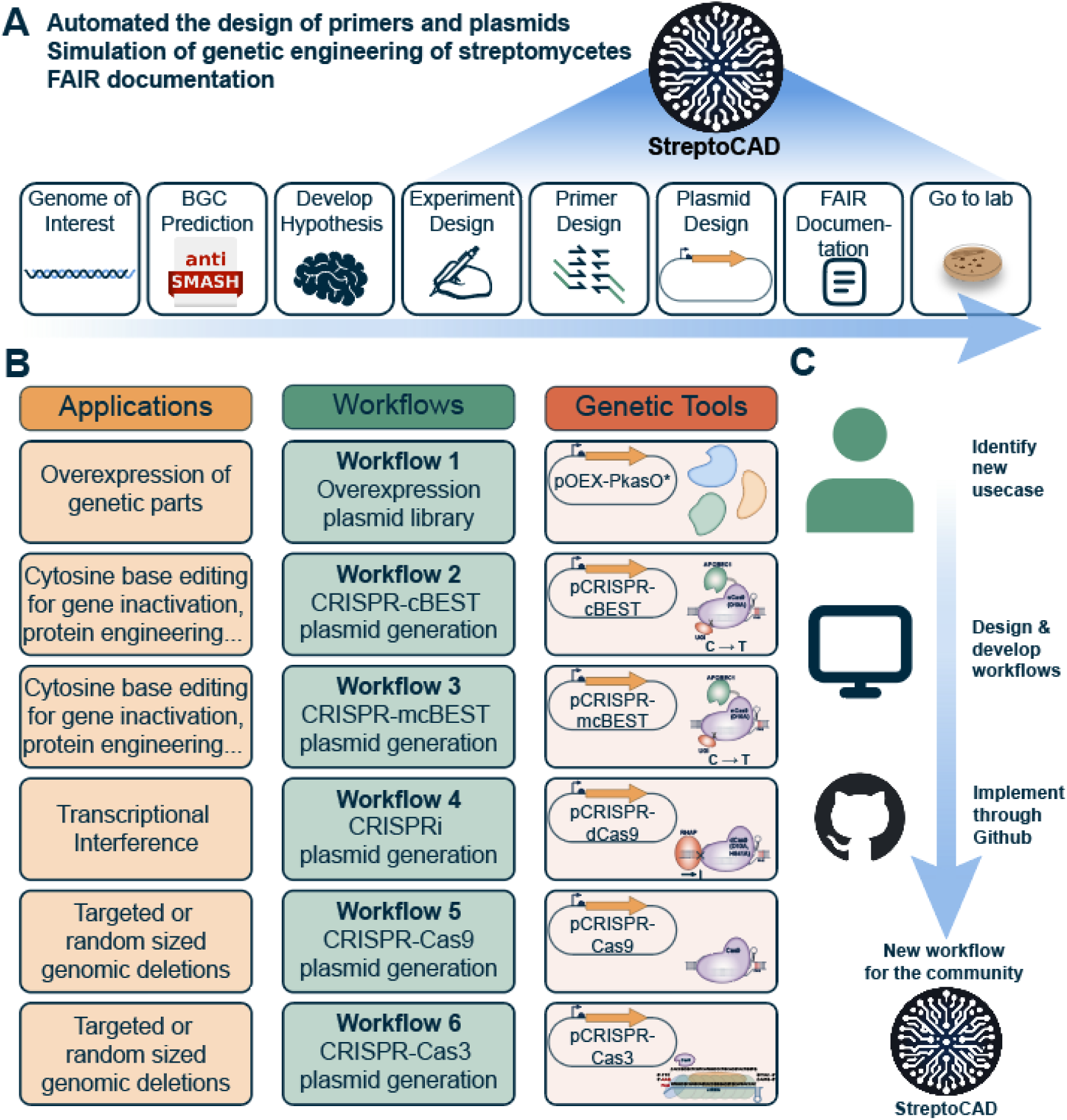
StreptoCAD features and community-driven efforts. **(A)** Generic workflow for *Streptomyces* genome engineering and highlights where StreptoCAD workflows can aid researchers, emphasizing how StreptoCAD fosters a shift from traditional methods to a more standardized, FAIR-compliant approach in bioengineering. **(B)** Summary of applications, workflows, and genetic tools for Streptomyces genome engineering. The figure outlines six workflows, from overexpression (Workflow 1) to CRISPR-based tools for base editing, transcriptional interference, and genomic deletions (Workflows 2–6), each linked to specific plasmids and genetic tools used to achieve these applications. **(C)** Community-driven development of workflows for genome engineering in *Streptomyces* through GitHub to benefit everyone.

### Design and Community Involvement

StreptoCAD is user-friendly, designed to be accessible to both novice and experienced users, with an intuitive interface and support for common biology file formats like GenBank and FASTA, facilitating easy integration with other tools and databases. As an open-source tool, it encourages community contributions, allowing researchers to add new workflows and customize the platform to meet specific research needs. Users can integrate new workflows by creating frontend files, developing callback functions, and writing tests, which can then be incorporated into the main application via GitHub (**Figure 1C**). Comprehensive documentation and user guides are available on GitHub (https://github.com/hiyama341/streptocad/blob/main/web_app/how_to_make_your_own_wor <u>kflows.md</u>), making it easier for users to contribute to and enhance the platform. This collaborative approach ensures that StreptoCAD evolves continuously, adapting to the latest advancements in genetic engineering and the diverse needs of the scientific community. Plasmids required for the different protocols are available at Addgene.

### Workflow 1: Gene overexpression plasmid library construction

Overexpression of potential activator genes is a common method to activate silent BGCs or to study the function of a given gene. Here, we developed a method to easily overexpress any gene using StreptoCAD. We demonstrate this by constructing a regulator overexpression library. This approach has proven effective in activating silent BGCs and enhancing the yields of natural products in *Streptomyces*^34–36^. To achieve this, we developed a rapid and standardized workflow for overexpressing regulators and constructed a plasmid system called pOEX-PkasO* (**Figure 2A, Suppl. Fig 7, Suppl. file 2**). This adapted plasmid allows for standardized cloning of any gene into the plasmid’s unique *StuI* restriction site, followed by PCR amplification of selected genes with defined overhangs, and subsequent Gibson assembly cloning. This system is based on the integrative plasmid pRM4e, described by Menges *et al.* 2007^37^, but incorporates one of the strongest known promoters engineered in *Streptomyces,* namely PkasO*^38^, along with the canonical RBS sequence “GGAGG”^39^. Additionally, the plasmid encodes the site-specific recombinase PhiC31 integrase, which mediates the integration of the plasmid’s contents into the *attB* site in chromosome^40^.

**Figure 2:**
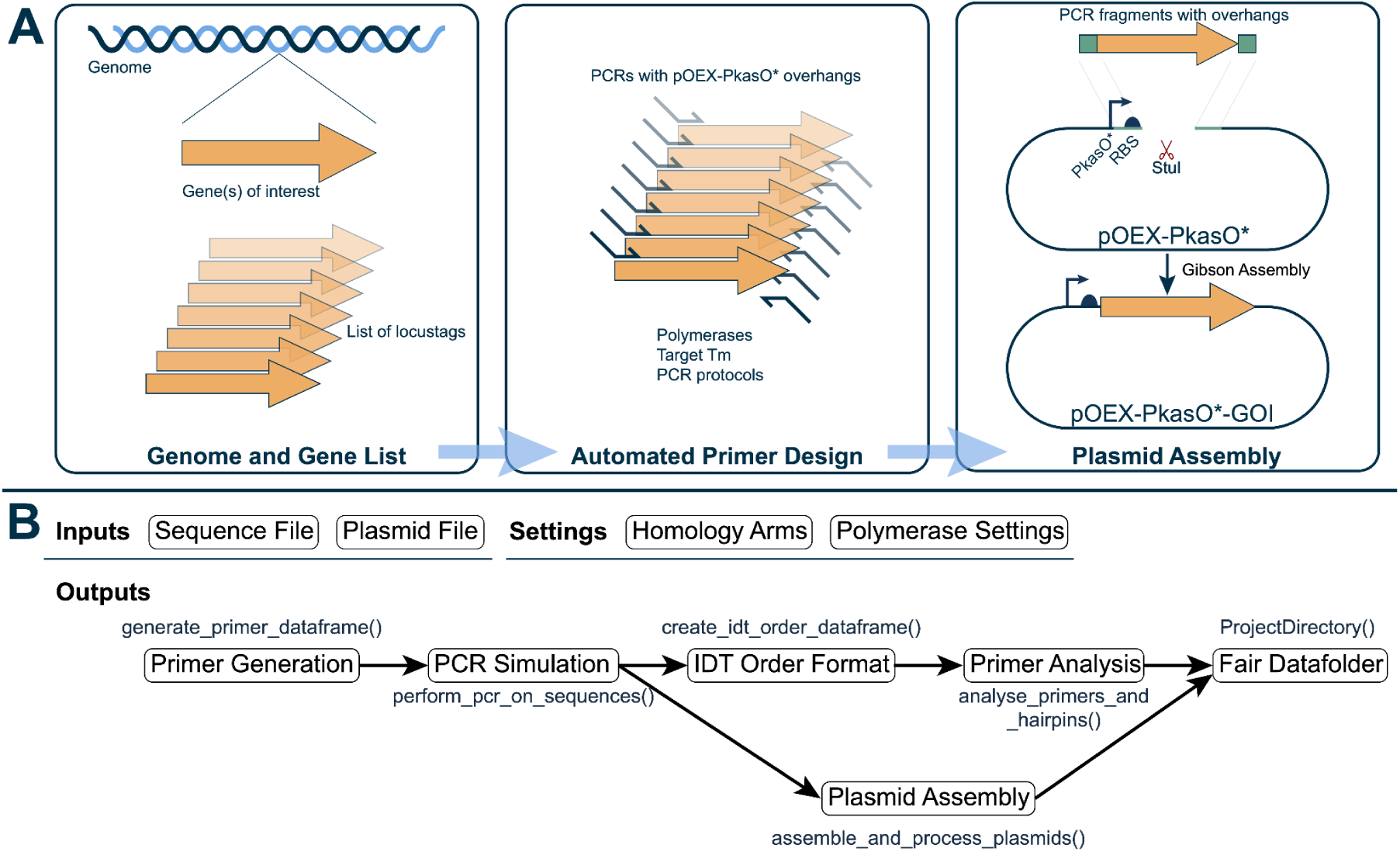
Overview of the Overexpression Workflow Using the pOEX-PkasO* Plasmid System. **(A)** Schematic of the overexpression process, starting from the identification of genes of interest (**left**) and automated primer design (**middle**) to plasmid assembly (**right**). The plasmid incorporates a unique *StuI* restriction site for cloning, a strong promoter (**P*kasO*\***) for high-level expression, and the canonical RBS sequence “GGAGG”. **(B)** T The workflow algorithm (W1-pOEX) illustrates the key algorithmic steps. The first step involves reading input files, which include sequences that the user wants to amplify, provided in a single GenBank file. Another required input is a plasmid file (pOEX-PkasO*), also in GenBank format. Example sequence and plasmid files can be downloaded using the respective download buttons. Additional settings, such as the polymerase and target melting temperature, can be modified. After reading the input files, the algorithm generates primers for each sequence with the specified overhangs using the generate_primer_dataframe function, which produces a DataFrame containing the primer sequences, anneal-specific sequences, melting temperatures, and annealing temperatures. The primers are validated through a PCR simulation using the perform_pcr_on_sequences function, which outputs a list of amplicons. Subsequently, the primers are compiled into a CSV order file using the create_idt_order_dataframe function, and plasmid assemblies are simulated with the assemble_and_process_plasmids function, which returns a list of contigs. Before the final step, the primers are analyzed for homo- and heterodimer formation as well as hairpins using the analyse_primers_and_hairpins function, which returns a DataFrame with the analysis. The final step produces a folder containing all input and output files used in the workflow, enabling reproducibility (Fair Data folder). Additional output files include all generated data as well as laboratory protocols necessary to perform the workflow in the lab..

StreptoCAD streamlines the overexpression design process by allowing users to input genes of interest in GenBank format, as shown in the "Inputs" section of **Figure 2B & Suppl. Fig 1.** StreptoCAD automatically generates primers with 5’-overhangs compatible with the pOEX-PkasO* plasmid (Forward primer overhang: 5’-GGCGAGCAACGGAGGTACGGACAGG*-gene-specific-sequence*-3’; Reverse primer overhang: 5’-CGCAAGCCGCCACTCGAACGGAAGG*-gene-specific-sequence*--5’), as illustrated in the "Primer Generation" and "PCR Simulation" stages. StreptoCAD then produces the corresponding plasmid constructs through automated assembly (depicted in the "Plasmid Assembly" step). **Figure 2B** also highlights outputs such as primer analysis and a comma-separated file containing primer names, sequences, and purification methods (IDT order format), alongside a downloadable data package that includes all inputs and outputs generated by StreptoCAD. This workflow has been experimentally validated, as described in the section “Experimental validation of workflow 1: Overexpression of transcriptional regulators in S. Gö40/10,” confirming its reliability in constructing overexpression plasmids.

### Workflow 2: CRISPR-cBEST plasmid generation for single edits

Workflow 2 focuses on the generation of plasmids for single-base edits using the CRISPR-BEST system, a double-strand break (DSB)-free approach that minimizes genomic instability and off-target effects. This workflow is built around the thermosensitive pSG5-based plasmid <u>CRISPR-cBEST</u>(Addgene: #125689)^13^ and the updated version <u>pCRISPR-cBEST-v2-PermE</u> (Addgene: #209446)^41^ for cytidine base editing.

The CRISPR-cBEST system utilizes a sgRNA to edit cytosines with single-nucleotide resolution, enabling precise introduction of point mutations, amino acid substitutions, or premature stop codons. The sgRNA cassette is driven by the constitutive P*ermE\** promoter, ensuring high transcription levels and efficient sgRNA expression (**Figure 3)**. The system’s core components include a *Streptomyces* codon-optimized cytidine deaminase (rAPOBEC1) fused to a Cas9 nickase (Cas9n D10A), and a uracil glycosylase inhibitor (UGI) to block uracil excision repair, thus facilitating efficient C-to-U conversion (**Figure 3A)**.

**Figure 3:**
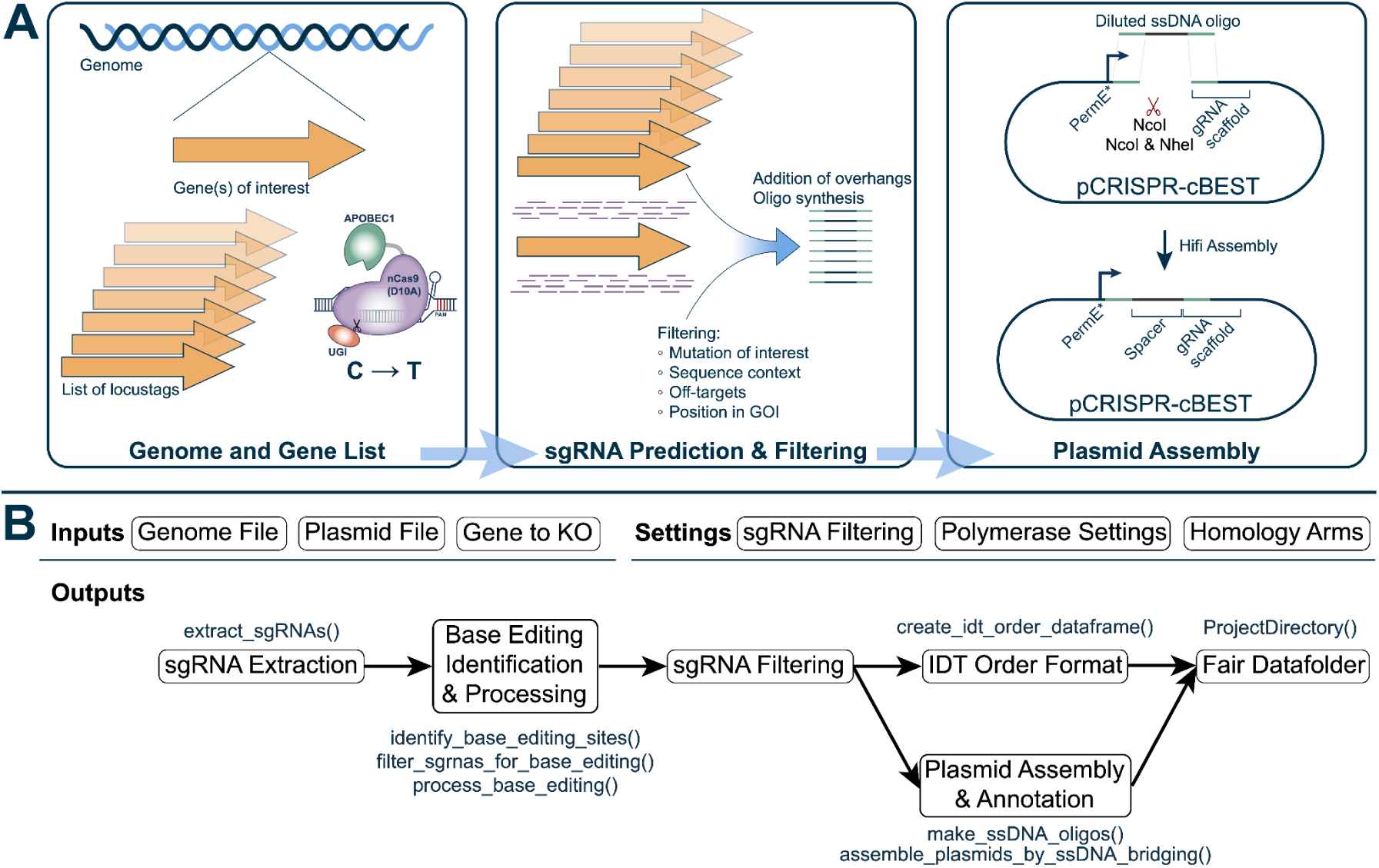
CRISPR-cBEST plasmid generation. **(A)**(**left**) Generating CRISPR-BEST sgRNAs from a list of locus tags or regions, (**middle**) filtering sgRNAs that are compatible with base-editing and making oligos. (**right**) Plasmid assembly simulation with the generated oligos and chosen restriction enzymes. **(B)** The W2-CRISPR-cBEST workflow algorithm illustrates the main computational steps. Inputs include genome and plasmid files, target genes or regions for base-editing, sgRNA filtering metrics, polymerase settings, and homology arms. The extract_sgRNAs function processes the genome and filtering metrics to identify all sgRNAs within the locus, outputting a DataFrame containing their location, strand, off-targets, and other relevant details. This is followed by three functions that identify the editing context, editable cytosines, and possible mutations based on base-editing, and then filter sgRNAs that do not meet the specified criteria (Base-editing Identification & Processing). Depending on the number of selected sgRNAs per locus tag or region, sgRNAs are filtered into a separate DataFrame (sgRNA-filtering). From this DataFrame, ssDNA oligos are designed using the makes_ssDNA_oligoes function, and plasmids are assembled via the assemble_plasmids_by_ssDNA_bridging function. Finally, sgRNAs are compiled into a CSV order file using the create_idt_order_dataframe function. All data, including inputs, outputs, and laboratory protocols, is stored in the final project folder to ensure reproducibility.

StreptoCAD streamlines the design of base-editing experiments by identifying suitable sgRNAs through its sgRNA identification tool, which locates and filters sgRNAs within a specified sequence based on their base-editing potential, as shown in **Figure 3B & Suppl. Fig 2**. The “Editing Sequence Context” feature further refines this selection by excluding sgRNAs with an upstream “G” adjacent to the editable “C,” ensuring higher editing efficiency (**Suppl. Fig 2)**. Additionally, when the "only stop codons" option is activated, StreptoCAD introduces an extra filtering step to pinpoint sgRNAs capable of introducing stop codons, as indicated in the sgRNA filtering stage of the figure.

As shown in **Figure 3B**, StreptoCAD proceeds to convert selected sgRNA sequences into oligos with 20 bp overhangs, enabling their integration into plasmids via single-stranded DNA (ssDNA) oligo bridging. This process, illustrated in the Plasmid Assembly & Annotation step, employs specific restriction sites (*NcoI*, or *NcoI & NheI*) and the ssDNA bridging method for seamless assembly. StreptoCAD automates the entire design workflow, generating plasmids, ssDNA bridging primers, and checking primers in IDT format, along with downloadable detailed protocols, as shown in the Outputs section of **Figure 3B**.

### Workflow 3: Multiplexed CRISPR-BEST plasmid generation

Workflow 3 leverages the <u>CRISPR-mcBEST</u> system (Addgene: #209416) for multiplexed genome editing, enabling the simultaneous targeted modification of up to 28 individual bases within a targeted genome^16^. Similar to <u>CRISPR-cBEST</u>, this workflow is used for cytidine deamination and is ideal for high-throughput genetic studies, bypassing the need for iterative strain engineering cycles.

The system uses multiple sgRNAs arranged as an array within a single vector, integrated using the Golden Gate cloning method. The Csy4/Cas6 system is used for processing the sgRNA array and releasing individual mature sgRNAs (**Figure 4A).** This plasmid has been optimized with the integration of the P*kasO*\* promoter to boost transcription levels of the sgRNA array and thereby editing efficiency (**Figure 4A)**. Correctly assembled plasmids are easily identified in this platform as the successful cloning of the sgRNA array into the plasmid results in the replacement of a sfGFP reporter gene, and the loss of fluorescence in *Escherichia coli*^16^.

**Figure 4:**
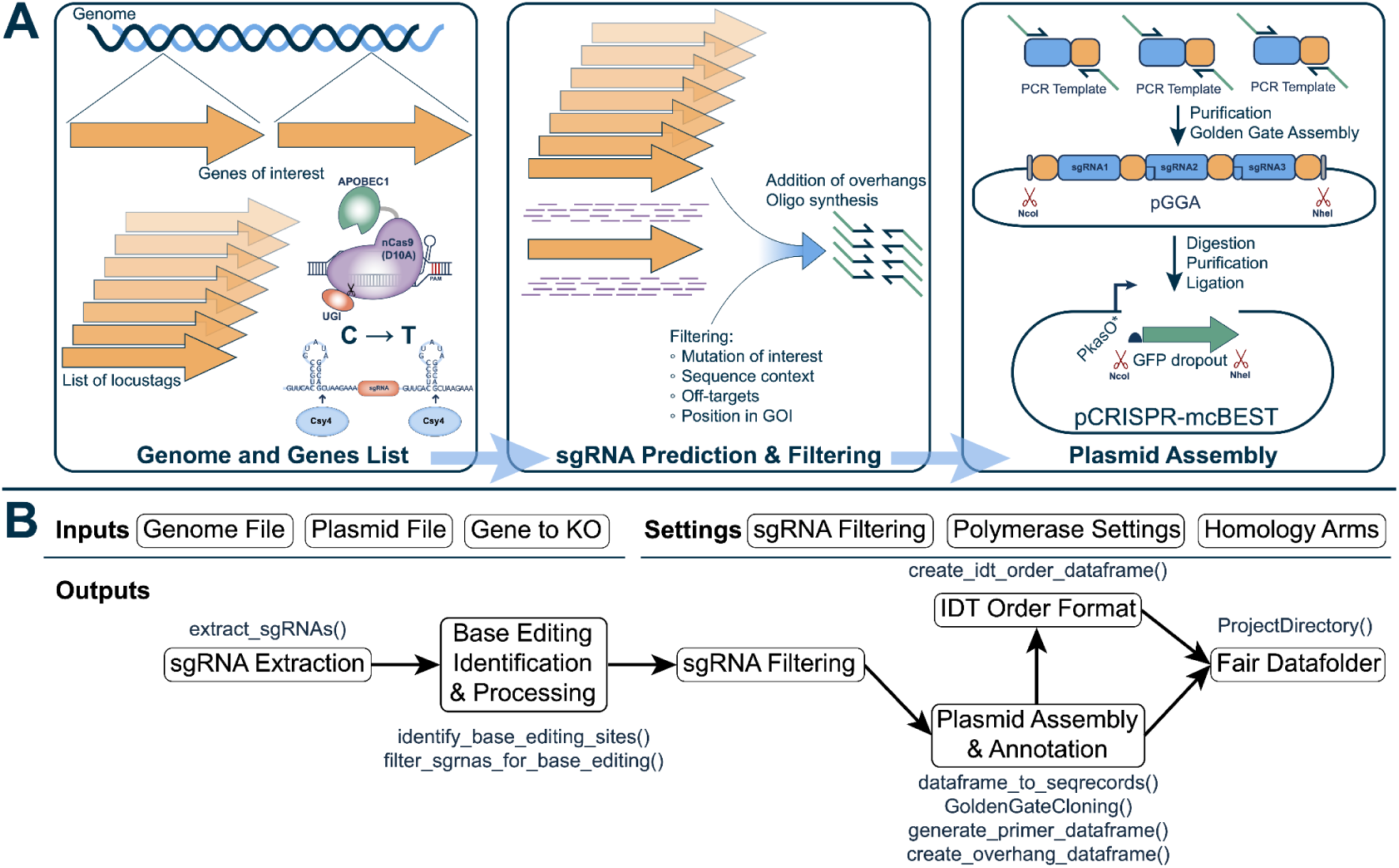
Generating a Multiplexed plasmid for base editing. **(A)** (**left**) Genes of interest are chosen for base-editing. (**middle**) sgRNAs are predicted and filtered based on criteria such as mutation of interest, sequence context, off-target potential, and their position within the gene of interest. Oligos are then generated with overhangs for downstream assembly. (**right**) A multiplexed plasmid is assembled using Golden Gate assembly, incorporating multiple sgRNAs and a GFP dropout as a reporter. The orange and blue PCR templates correspond to the sgRNA handle containing the Cys4 sequence. Restriction enzymes (e.g., NcoI, NheI) facilitate plasmid construction through digestion, purification, and ligation. **(B)** The W3-CRISPR-mcBEST workflow algorithm shares similarities with the W2-CRISPR-cBEST workflow but utilizes Golden Gate assembly for plasmid construction and supports multiplexing. Inputs include genome and plasmid files, target genes for knockout (KO), sgRNA filtering metrics, polymerase settings, and homology arms. The workflow begins with sgRNA extraction using the extract_sgRNAs function, which identifies all sgRNAs within the specified locus based on the filtering metrics and outputs a DataFrame containing sgRNA locations, strands, and off-target information. Next, the editing context, editable cytosines, and possible mutations are identified, and sgRNAs are filtered to meet the criteria for base-editing identification and processing. From the filtered sgRNAs, the dataframe_to_seqrecords function converts the data into sequence records for downstream applications. Golden Gate assembly is performed using the GoldenGateCloning class, which automates the creation of multiplexed plasmid assemblies. Primers are generated using the generate_primer_dataframe function, while overhang sequences are designed with the create_overhang_dataframe function. The final outputs include a CSV order file in IDT order format and a FAIR data folder containing all inputs, outputs, and protocols necessary for reproducibility and laboratory execution.

StreptoCAD streamlines sgRNA selection and processing by identifying sgRNAs with low off-target hits. It then generates primers for amplifying individual sgRNAs from the sgRNA handle containing Cys4, including overhangs for directional assembly into the plasmid, as shown in **Figure 4B & Suppl. Fig 3**. To prevent issues with duplicate overhangs across multiple individual sgRNAs, StreptoCAD generates a table highlighting all overhangs and their potential duplications, ensuring sequence uniqueness within the sgRNA array. The software also simulates the Golden Gate purification process, including *BsaI* digestion of the PCR fragments and the circular assembly of the plasmid, as illustrated in the Plasmid Assembly step (**Figure 4B**).

As in workflow 2, the functions “Editing Sequence Context” and “Only Stop Codons” refine sgRNA selection for targeted base-editing applications. Unlike the other workflows, Workflow 3 supports multiplexing by generating a single plasmid capable of housing multiple sgRNAs. However, users must submit separate requests if multiple plasmids are required, adding flexibility for more complex experiments.

### Workflow 4: CRISPRi plasmid generation

Workflow 4 uses CRISPR interference (CRISPRi) to achieve targeted gene silencing in bacteria. This system utilizes a dead-Cas9 (dCas9) protein complexed with a guide RNA (gRNA), which binds to specific genomic regions and blocks RNA polymerase, effectively halting transcription. The CRISPRi system used in this workflow is based on the <u>pCRISPR-dCas9</u> plasmid (Addgene: #125687), which was constructed by mutating both nuclease domains (D10A and H840A).

The sgRNA scaffold is expressed under the constitutive P*ermE*\* promoter. A single *NcoI* restriction site flanks the 20-nucleotide target sequence within the sgRNA, allowing for easy insertion of specific targets (**Figure 5A)**. Expression of the dCas9 encoding gene is regulated by the thiostrepton-inducible P*tipA* promoter^33^.

**Figure 5:**
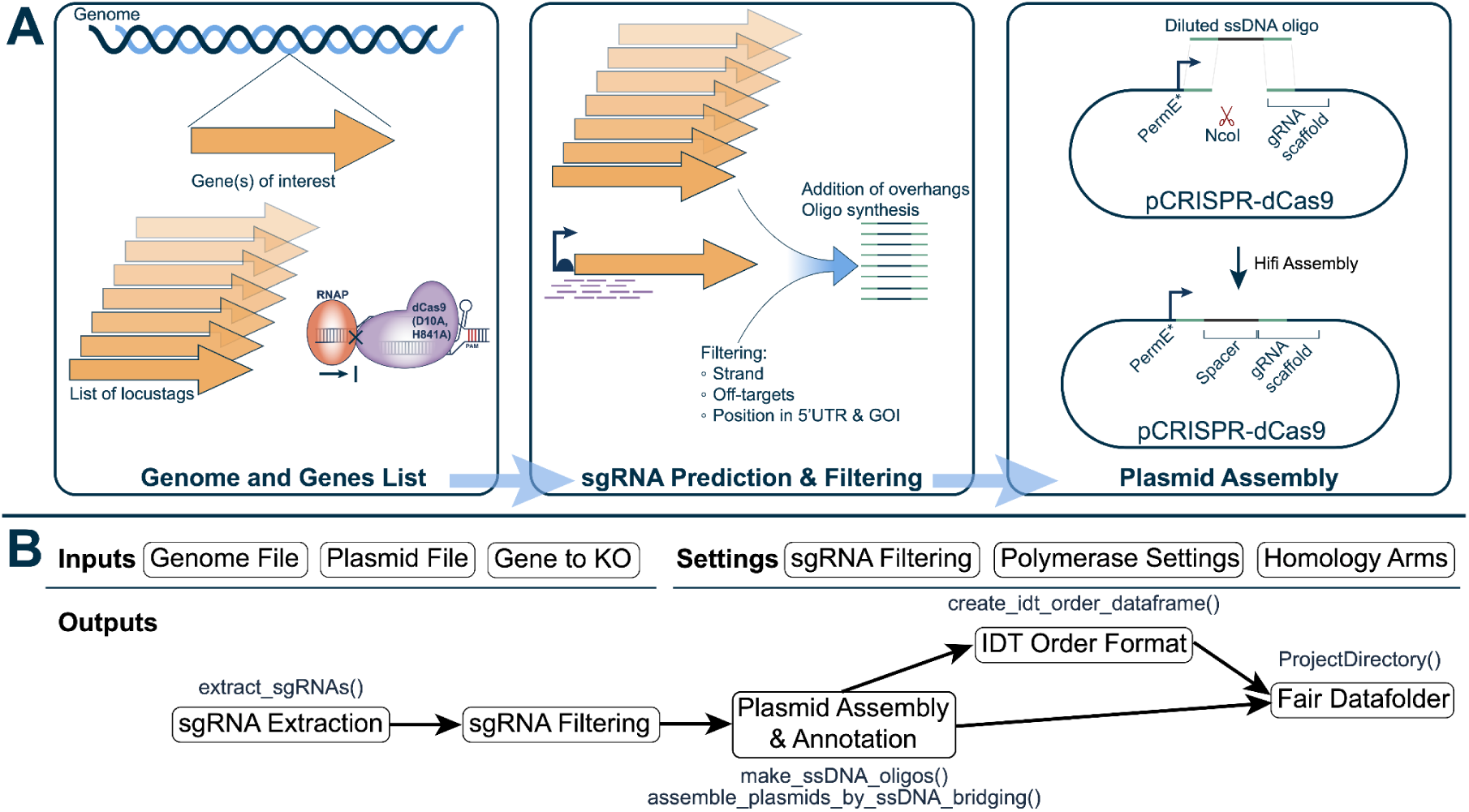
Generating single sgRNA CRISPRi plasmids. **(A)** (**left**) A list of locus tags corresponding to target genes is used to guide sgRNA design. The CRISPRi mechanism, relying on transcriptional inhibition by dCas9, is illustrated. (**middle**) sgRNAs are identified from the regions upstream and around the transcription start site (TSS) of the designated locus tags. These sgRNAs are sorted, filtered based on strand, off-target potential, and proximity to the TSS, and oligos are generated with the appropriate overhangs. (**right**) Simulation of plasmid assembly is performed using the synthesized oligos, generating a CRISPRi plasmid with the required sgRNA scaffold and regulatory elements. **(B)** The W4-CRISPRi workflow algorithm illustrates the key computational steps. Inputs include genome and plasmid files, target genes for transcriptional inhibition, sgRNA filtering metrics, polymerase settings, and homology arms. The workflow begins with sgRNA extraction using the extract_sgRNAs_for_crispri function, which finds sgRNAs upstream of the transcription start site (TSS) and outputs a sorted DataFrame containing sgRNA locations ranked by proximity to the user-defined upstream value. Next, the user-specified number of sgRNAs are selected and filtered (sgRNA Filtering). Filtered sgRNAs are then used to design ssDNA oligos with the make_ssDNA_oligoes function and to assemble plasmids through the assemble_plasmids_by_ssDNA_bridging function. Primers are generated and compiled into a CSV file in IDT order format using the create_idt_order_dataframe function. Finally, all input and output data, including laboratory protocols required to execute the workflow, are compiled into a downloadable folder for reproducibility.

CRISPRi can inhibit transcription either by blocking transcription initiation or elongation, depending on the targeted region^42^. To block transcription initiation, CRISPRi targets promoter regions located upstream of the transcriptional start site (TSS) on either the template or non-template strand. In prokaryotes, promoters are typically found within 200 bp upstream. These regions are crucial for RNA polymerase binding, making promoter targeting an effective strategy for gene silencing^43^. When blocking transcription elongation, CRISPRi is only effective when targeting the non-template strand^42^.

StreptoCAD streamlines the CRISPRi design process by automatically selecting the appropriate non-template strand and allowing users to define a region upstream of the transcription start site (TSS), with the default set to 100 bp to accommodate most bacterial promoters, as depicted in **Figure 5B & Suppl. Fig 4**. The software simulates plasmid assembly by digesting the input plasmid with *NcoI* and incorporating sgRNAs into single-stranded DNAs (ssDNAs), as shown in the Plasmid Assembly stage. Standardized overhangs (5’-CGGTTGGTAGGATCGACGGC-**N20**-GTTTTAGAGCTAGAAATAGC-3’) are added to the sgRNA, resulting in a 60mer used for ssDNA bridging (**Figure 5B**).

Since CRISPRi does not involve permanent genetic modifications, StreptoCAD omits the generation of checking primers in this workflow. Instead, efficiency validation requires transcriptomic analysis or endpoint product-level assessment. As with other workflows, users can download a comprehensive data package, including all inputs, outputs, and protocols, by selecting "Download all data & protocols" (**Figure 5B**).

### Workflow 5: Deletion using CRISPR-Cas9

Workflow 5 utilizes the well-established CRISPR-Cas9 system to introduce DSBs for generating random-sized or in-frame deletions, depending on whether an editing template is provided (**Figure 6A**).

The <u>pCRISPR-Cas9</u> system (Addgene: #125686), developed by Tong *et al.* 2015^33^, utilizes a plasmid backbone from pGM1190, which includes the temperature-sensitive replication origin pSG5^44^ and an apramycin resistance marker. The temperature sensitivity of the vector facilitates the easy removal of plasmids by raising the incubation temperature above 37°C, ensuring that introduced genetic changes remain stable without the continuous presence of the plasmid. Cas9 expression is regulated by the thiostrepton-inducible P*tipA*\* promoter^45^. The sgRNA cassette is expressed from the constitutive P*ermE*\* promoter, for the constitutive expression of sgRNAs.

As shown in **Figure 6B**, StreptoCAD streamlines the design of the assembly process by filtering and selecting sgRNAs and simulating their incorporation into plasmids via ssDNA oligo bridging. The workflow includes digesting the plasmid with *NcoI* and performing *in silico* assembly. When the "Generate in-frame deletions" option is enabled, StreptoCAD designs a repair template flanking the target gene or region with 1 kb upstream and downstream sequences. Primers with overlapping regions for Gibson cloning are generated, and the second plasmid assembly step is simulated by digesting the sgRNA-integrated plasmid with *StuI* or any of the enzymes from the MCS. The software then performs a Gibson cloning simulation to validate the plasmid design, ensuring seamless assembly. The process concludes with the generation of an IDT order format and a FAIR data package for download (**Suppl. Fig 5**).

### Workflow 6: Deletion using CRISPR-Cas3

Workflow 6 utilizes the <u>pCRISPR-Cas3</u> plasmid (Addgene: #209427), based on the compact type I-C CRISPR system developed by Csörgő et al.^46^, to introduce deletions or substitutions designed for *Streptomyces* species.^18^. The CASCADE-Cas3 system employs a multi-effector CASCADE complex expressed from a single P*tipA* promoter in a polycistronic operon consisting of Cas5, Cas8, Cas7, and Cas3. This CASCADE complex recognizes the target DNA and recruits Cas3 to initiate DNA degradation. Like other CRISPR plasmids in this study, pCRISPR-Cas3 utilizes the temperature-sensitive pSG5^44^ replicon and includes apramycin resistance as a selectable marker.

pCRISPR-Cas3 is particularly effective for large deletions (>100 kb), such as removing entire biosynthetic gene clusters (BGCs), but is also suitable for small (∼ 0.1-7 kb) and mid-sized deletions (∼7-100 kb), as well as random-sized deletions^46^. It can replace large genomic regions with specific cargo sequences or repair templates, achieving high efficiencies for targeted deletions. Cas3 recognizes a T-rich 5’-TTC-3’ protospacer adjacent motif (PAM), which reduces off-target hits in high GC organisms like *Streptomyces*. The system uses a longer 34-nt protospacer, with expression driven by the constitutive P*ermE\** promoter.

StreptoCAD automates the full pCRISPR-Cas3 design workflow, simulating each step from plasmid construction to final assembly (**Figure 7A & 7B)**. The workflow begins by selecting crRNAs based on user-defined filtering metrics. Next, primers are generated with crRNAs incorporated into the primer overhangs along with two fixed primer sequences (Forward primer: 5‘-GAGCTCATAAGTTCCTATTCCGAAG-3’; Reverse primer: 5‘-AAGAAGTGGGTGTCGGACG-3’) for PCR amplification (**Figure 7A & 7B**). StreptoCAD then simulates plasmid assembly that starts with plasmid digestion using *BstBI* and *NdeI* followed by Gibson cloning with the amplified crRNAs incorporated into the insert, accounting for the high GC content and sequence homologies typical of *Streptomyces* species (**Figure 7A**).

**Figure 6:**
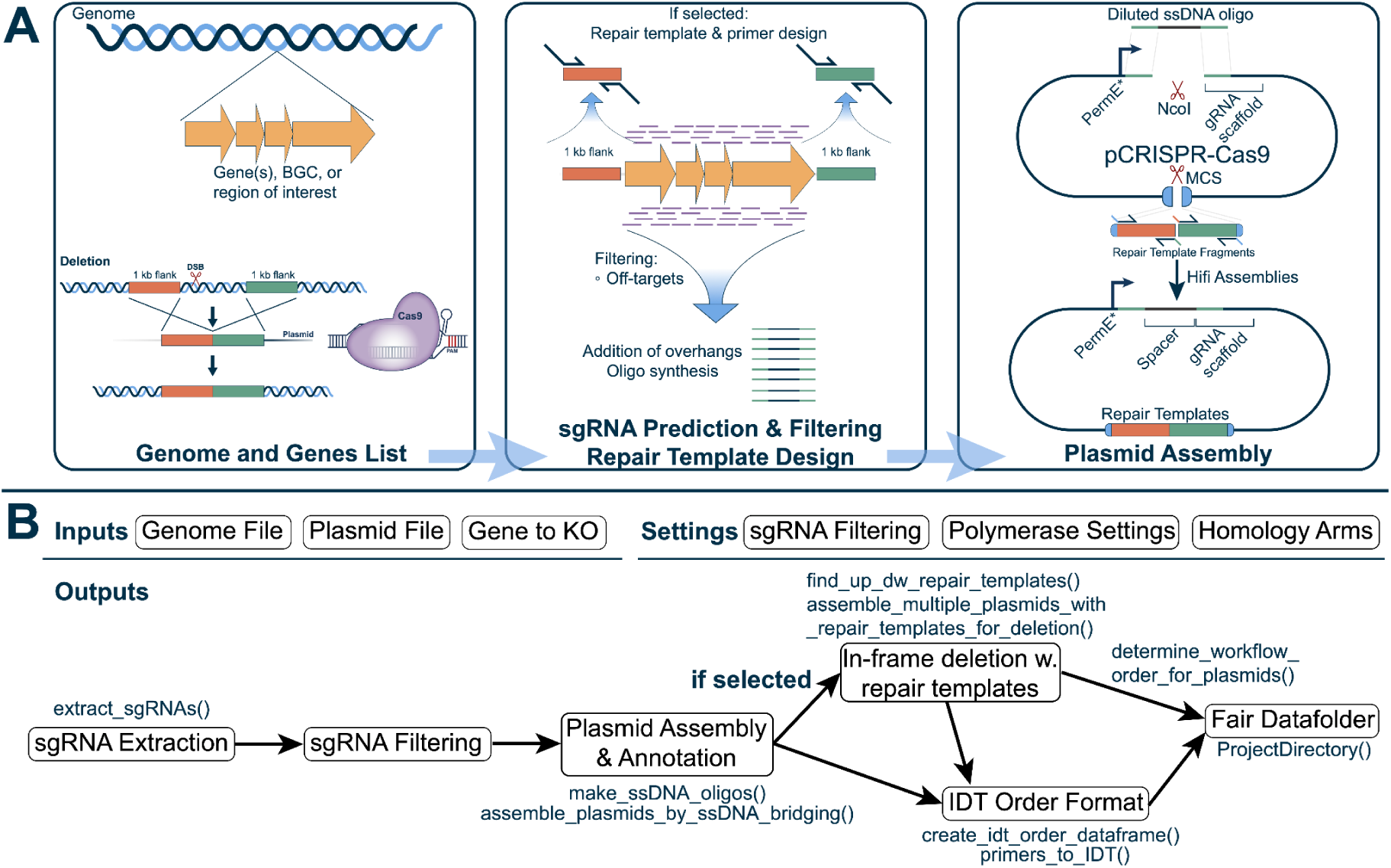
Generation CRISPR-Cas9 plasmids. **(A)**(**left**) Genome and locus tags/regions are parsed and (**middle**) sgRNAs are predicted, filtered and oligos made. If an in-frame deletion is chosen, 1kb fragments are found upstream and downstream of the locus tags/regions and oligos to amplify these are made with overhangs to perform Gibson assembly. (**right**) CRISPR-Cas9 plasmid assembly simulation is simulated in two steps, i) ssDNA bridging with the sgRNA oligo is simulated ii) Gibson cloning with upstream and downstream repair templates is simulated. (**B**) The W5-CRISPR-Cas9 In-Frame Deletion workflow algorithm illustrates the key computational steps. Inputs include genome and plasmid files, the target gene or region, sgRNA filtering metrics, polymerase settings, and homology arms. The workflow begins with sgRNA extraction using the extract_sgRNAs function, which finds sgRNAs in the target region. These sgRNAs are filtered according to user-defined parameters. The sgRNA oligos are then generated by the make_ssDNA_oligoes function and assembled into a CRISPR-Cas9 plasmid using the assemble_plasmids_by_ssDNA_bridging function. If in-frame deletion is selected, the find_up_dw_repair_templates function identifies 1 kb upstream and downstream fragments, and oligos overlapping eachotheer are made for the HifI assembly. with the assemble_multiple_plasmids_with_repair_templates_for_deletion function. Final outputs include a CSV order file in IDT order format with all oligoes as well as validation primers for the in-frame deletion, a FAIR data folder containing all input and output files, and laboratory protocols necessary to reproduce the workflow.

**Figure 7:**
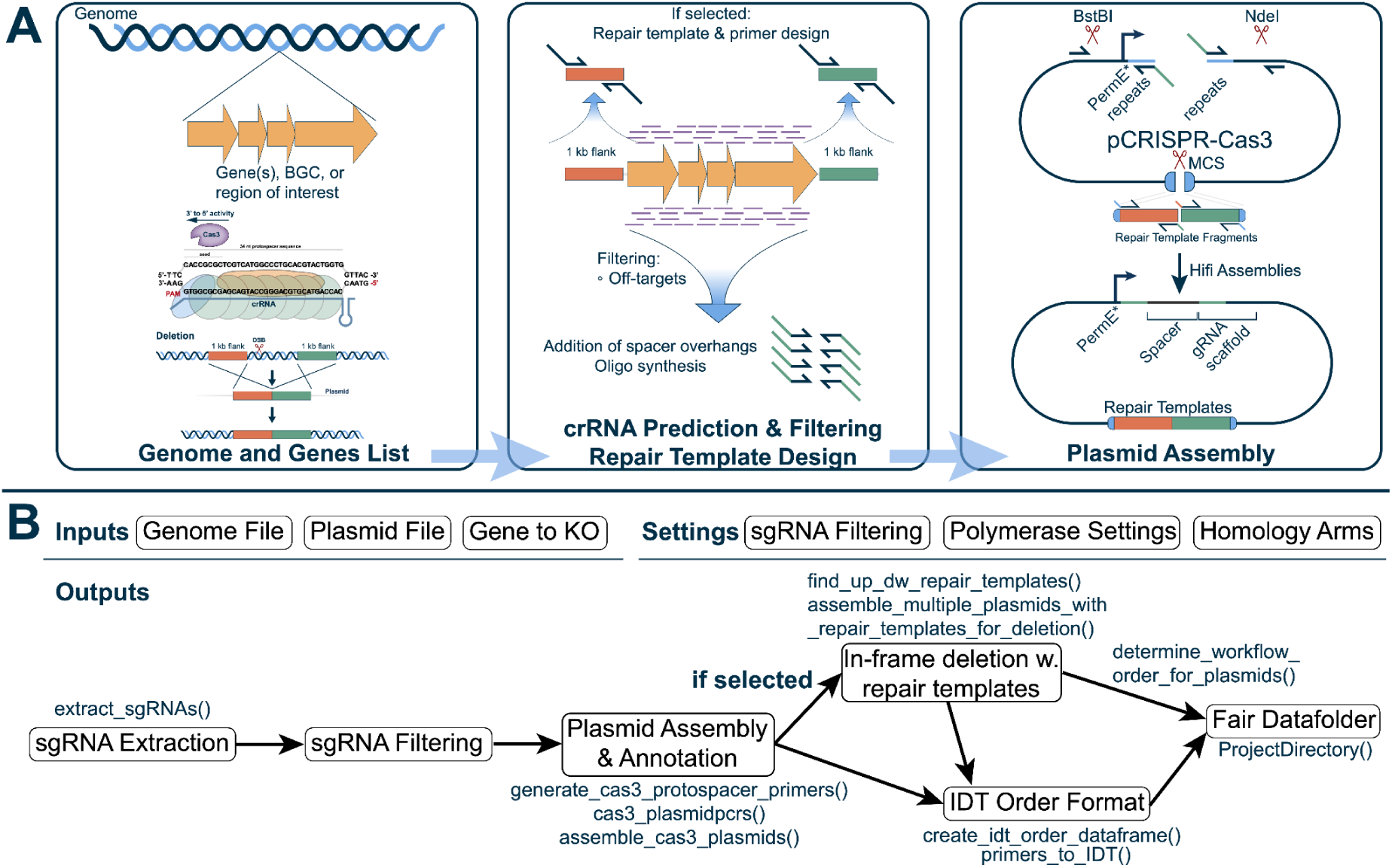
Generation CRISPR-Cas3 plasmids. **(A)**(**left**) Locus tags or regions are chosen and (**middle**) crRNAs are found and filtered. Oligos for integrating the crRNAs through Gibson assembly are generated. If the selected repair templates are also generated with primers compatible with Gibson assembly. (**right**) integrating crRNAs through PCR amplification and subsequent Gibson assembly. If repair templates are selected these are included in the assembly simulation process. (**B**) The workflow begins with crRNA extraction using the extract_sgRNAs function. These crRNAs are filtered using user-defined parameters (e.g., strand, off-target scores). The filtered crRNAs are used to design oligos via the generate_cas3_protospacer_primers function and are assembled into plasmids using the cas3_plasmidpcrs and assemble_cas3_plasmids functions. If repair templates are selected, the find_up_dw_repair_templates function identifies upstream and downstream fragments, and primers overlapping these fragments are generated. The plasmid is then assembled through Hifi assembly using the assemble_multiple_plasmids_with_repair_templates_for_deletion function. Final outputs include a CSV order file in IDT order format, containing all crRNA oligos and repair template primers in a FAIR data folder with all input and output file and laboratory protocols necessary for executing the workflow, ensuring reproducibility.

StreptoCAD further allows the user to design full in-frame deletions (removal of entire ORF) by activating the "Generate in-frame deletion" option.In this case, StreptoCAD proceeds with a second digestion and assembly step. Users can choose their restriction enzymes for the multiple cloning site (default: *EcoRI*) (**Figure 7A & B**). Subsequently, 1 kb upstream and downstream sequences of the target gene or region are retrieved and primers are designed to amplify and incorporate these into the plasmid via Gibson cloning (**Figure 7B**). The final plasmid design is validated through *in silico* assembly. As with other workflows, a comprehensive data package is generated, including the IDT order format, repair template sequences, and assembly protocols, ready for experimental use (**Figure 7B**, **Suppl. Fig 6**).

### Experimental validation of Workflow 1: Overexpression of transcriptional regulators in *Streptomyces* sp. Gö40/10 *produces novel phenotypes*

To demonstrate how StreptoCAD can simplify and streamline engineering workflows by automating the design step, we generated a transcriptional regulator overexpression library using Workflow 1. This genetic engineering strategy has been described by <u>Beck *et al.* 2021</u>, who successfully identified and overexpressed a set of transcriptional regulators from the SARP family. Analysis of the resulting genetically engineered strains allowed Beck *et al*. to identify and activate the griseusin cluster in *Streptomyces* sp. CA-256286^47^. Inspired by this work, StreptoCAD was used to construct plasmids for overexpression of a series of putative regulators, potentially resulting in the activation of silent BGCs.

To this end, we engineered a novel plasmid based on pRM4^37^ (**Figure 2A**), incorporating a standardized integration site with a *StuI* restriction site downstream of the strong promoter *PkasO\** and the canonical ribosome binding site sequence (5’-**GGAGG-3’**). We selected to work with *Streptomyces* sp. Gö40/10 (*Streptomyces* Gö40/10), a strain previously described as a producer of collinolactone, a potential anti-alzheimers drug^47–49^. Additionally, *Streptomyces* Gö40/10 is predicted to harbor 39 BGCs, of which the majority remains uncharacterized (**Suppl. File 2**). Thus, *Streptomyces* Gö40/10 serves as a good test bed for the workflow for activation of BGCs by overexpression of transcriptional regulators.

For the design of our library, LuxR and SARP regulators were selected. These regulator families frequently act as activators and have previously been shown to activate BGCs^36^. The antiSMASH output was parsed to identify all LuxR and SARP family regulators present in the *Streptomyces* Gö40/10 genome sequence (<u>00-parsing_antiSMASH.ipynb</u>). Resulting in the identification of 18 putative regulators located within 13 predicted BGCs, 7 from the SARP family and 11 from the LuxR family of transcriptional regulators (**S1 table 1**). Using the default settings of StreptoCAD, the 18 regulator sequences were uploaded in GenBank format along with the **pOEX-PkasO\*** plasmid sequence. StreptoCAD facilitated the design of all necessary primers for amplification of the regulators including 5’-overhangs with homology regions to allow for directed incorporation into the *StuI* site of the **pOEX-PkasO\*** vector (**Figure 2F**). A detailed description of this work is available in Supplementary material (**<u>01-Construction_of_pOEX_PkasO_capture_plasmid.ipynb</u> and <u>02-Integration_of_GÖ4010_regulators_into_pOEX_PkasO.ipynb</u>)**.

In the laboratory, the gDNA was isolated and PCR amplifications were performed on the 18 targeted regulators, of which 17 were successfully amplified. The purified PCR products were cloned into the *StuI* digested pOEX-PkasO* vector and subsequent transformations into *E. coli* yielded 15 correctly assembled plasmids, as confirmed by Sanger sequencing. These results were achieved without troubleshooting steps, suggesting that additional refinement could potentially yield the complete set of constructs. The validated plasmids were then transformed into the *E. coli* ET12567 pUZ8002 strain. The successfully transformed *E. coli* ET12567 puZ8002 strains were then used for conjugative transfer of the regulator overexpression library into *Streptomyces* Gö40/10. Conjugations resulted in 13 successful plasmid transfers and integrations, as indicated by antibiotic-resistant *Streptomyces* Gö40/10 colonies and validation by diagnostic colony-PCR using the primers designed by StreptoCAD (**S2 table 2, <u>02-Integration_of_GÖ4010_regulators_into_pOEX_PkasO.ipynb</u>**, **Suppl. Figure 8**).

To evaluate the phenotypical impact of overexpressing the individual 13 regulators on the strains’ phenotype, we inoculated the individual strains on DPNM medium **Figure 8B and Figure 8C**. The culturing revealed phenotypic variations among the generated strains, confirming successful genetic modifications through the integration of LUXR and SARP regulators. Most notably, the strain modified with Regulator 14, a LuxR family DNA-binding response regulator, located in BGC 28, (**Fig. 8**; **S2 table 2**) showed a noticeable change of color to an orange hue. AntiSMASH predicts this compound to be a polyketide and the BGC shows 89 % similarity to the A-74528 biosynthetic gene cluster (**Suppl. File 2**) from *Streptomyces* sp. SANK 61196 (GenBank: GU937384.1), which produces a C_30_ polyketide^50^. This distinct phenotypic alteration warrants further investigation to elucidate the specific compounds produced by this strain, which could potentially be a novel bioactive molecule. Additionally, the strain carrying Regulator 18 demonstrated a unique phenotype characterized by a darker hue at the center of the colony, similar to observations in the strain with Regulator 4. These color variations suggest alterations in metabolic pathways that are significant enough to affect pigment production.

**Figure 8:**
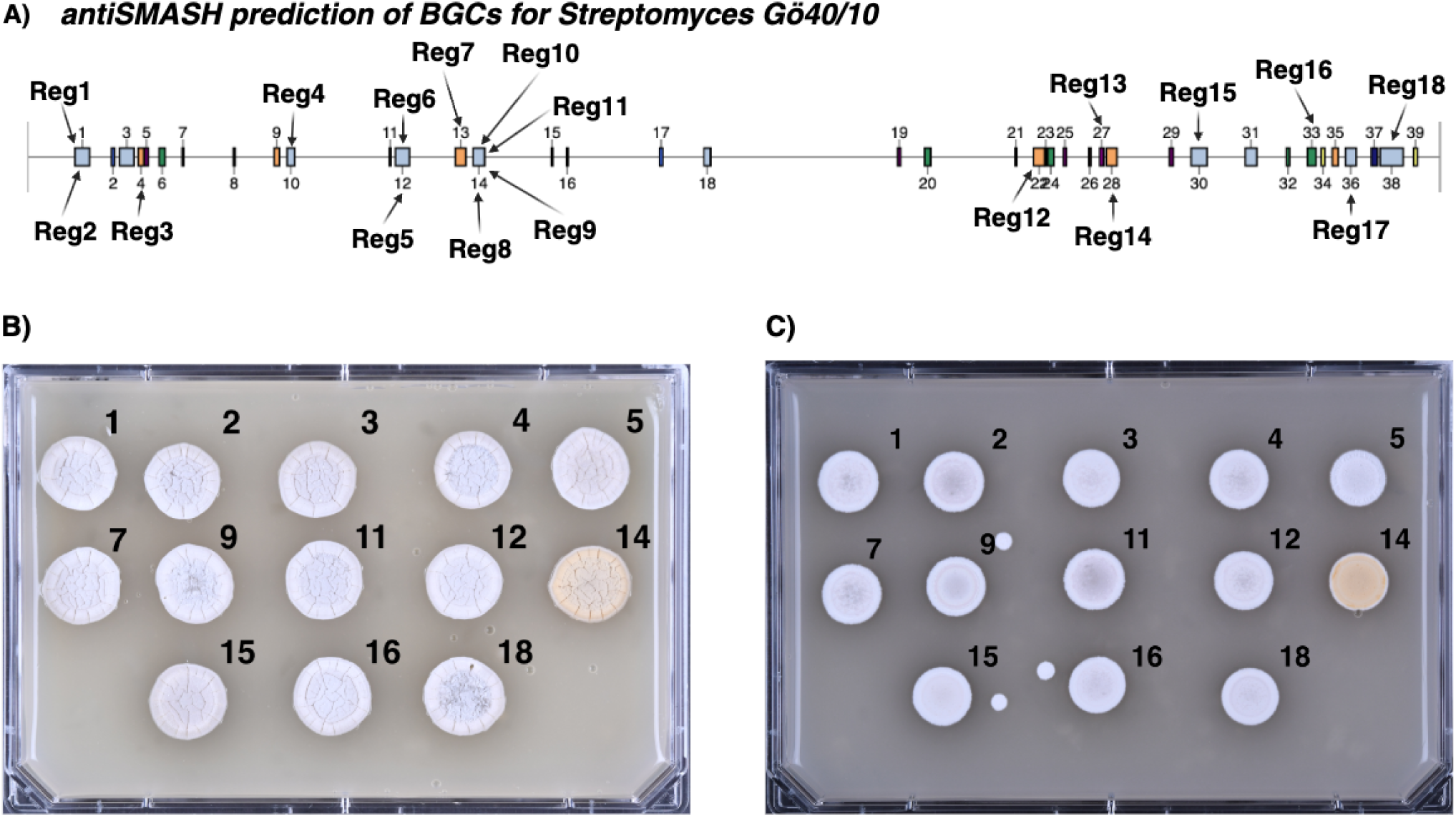
Location of overexpressed BGC regulators and the phenotypic effect of overexpression. **(A)** AntiSMASH prediction of BGCs in the *Streptomyces* Gö40/10 genome with the 18 selected regulators (Reg1-18) highlighted. (**B**) The 13 successfully constructed overexpression strains cultured on DNPM media supplemented with Apramycin (**C**) and on MS media supplemented with Apramycin. The WT strain was not included as it does not grow on antibiotic-containing plates.

The combined data show that the genetic engineering strategy and materials designed by StreptoCAD were functional and allowed for efficient implementation.

## Discussion

Designing genome engineering strategies is often a tedious task with poor reproducibility due to inconsistent design guidelines across designers/laboratories. While laboratory automation has accelerated the practical aspects of strain engineering, the design process is still predominantly manual. Therefore, this study showcases StreptoCAD as a design tool for i) automating and increasing the speed of genome engineering of *Streptomyces*, ii) standardizing the design process to minimize errors and ensure consistent cross-experiment behavior, iii) improving reproducibility by adhering to the FAIR principles by fully documenting all experimental design decisions, and iv) autogenerating all the experiment-specific laboratory protocols needed for implementation in the laboratory We have demonstrated that StreptoCAD can streamline experimental workflows, which we experimentally validated by constructing an overexpression library of transcriptional regulators. StreptoCAD enabled a beginner in *Streptomyces* engineering to generate and validate new strains in just eight weeks. The design phase, typically performed manually and taking several hours per strain, was completed in less than one minute after identifying the target genes for analysis. Although we achieved a success rate of 70%, this reflects our deliberate focus on rapid construction without repeating experimental steps or performing troubleshooting, rather than any limitation of the design tool. Specifically, out of the 18 targeted regulators, one failed during PCR amplification, two failed during plasmid assembly, and two grew slower than the rest in the *E. coli* ET12567 puZ8002 strain and were therefore skipped. These failures are typical issues that can be resolved through iterative optimization. By intentionally omitting troubleshooting to prioritize speed, we demonstrated StreptoCAD’s capability to deliver swift results even under these constraints. StreptoCAD’s speed highlights that the design process is no longer a bottleneck, as we have effectively solved this part of the DBTL cycle. The primary constraints now lie in the manual labor and the number of designs we can feasibly manage, demonstrating StreptoCAD’s potential for significantly scaling up strain construction workflows. This rapid generation process underscores StreptoCAD’s user-friendly nature and efficiency, allowing researchers to progress quickly through workflows without significant troubleshooting. Moreover, as the number of designs increases, the time savings become even more substantial. Future simulations could further evaluate StreptoCAD’s scalability and efficiency in handling larger-scale designs.

While several tools are available for designing molecular biology and genetic engineering strategies, StreptoCAD is unique in its focus on full CRISPR workflows and for the overexpression of any gene of interest in *Streptomyces*. StreptoCAD fills this niche and can potentially integrate functionalities from other platforms, such as the combinatorial mismatch of base pairs from CRISPy-web, to enhance its CRISPR-related capabilities^28^. The underlying coding/scripting design of StreptoCAD is highly modular, allowing for the reuse of elements/functions and code that can be combined in various ways. This modularity enables StreptoCAD to be adapted for use with other organisms and expanded with new modules, such as those for different CRISPR variants or additional gene editing tools, making it a versatile platform for diverse genetic engineering applications.

StreptoCAD offers significant out-of-the-box functionality that complements other available tools. For example, AutoESDCas^51^ is a powerful tool for whole-workflow editing sequence design using CRISPR/Cas systems and supports various genetic manipulations and high-throughput strain modifications. Similarly, ClusterCAD aids in the design and engineering of PKS and NRPS clusters by providing a database and tools for combinatorial biosynthesis based on sequence and structural similarity^52^. Galaxy-SynBioCAD^23^ provides robust capabilities for synthetic biology, metabolic engineering, and industrial biotechnology, supporting the entire pathway design process and the creation of strain libraries. TeselaGen and Geneious also provide comprehensive tools for DNA design, assembly, and bioinformatics^22,53^. However, these platforms often require users to go through extensive tutorials to utilize their full potential. In contrast, StreptoCAD is designed to be intuitive and user-friendly, enabling researchers to quickly adopt and apply its features without the need for extensive training.

Current limitations of StreptoCAD include a finite number of available workflows and the potential complexity of creating new workflows, particularly for users without a background in Python programming. While the web app format of StreptoCAD offers great accessibility, it may limit flexibility, which could be mitigated by integrating a Jupyter-based environment for advanced users. Importantly, StreptoCAD’s inherent ability to standardize designs will be crucial for conducting meta-studies that combine results from multiple experiments and groups. This standardization will allow for a more reliable comparison of outcomes across different studies, based on the standardized design rules, thereby enhancing our understanding of when and why certain genetic engineering approaches fail and improving overall scientific practice. This approach can “close the loop” by providing insights into underlying biological processes, making the design process more effective over time. By automating the design process with defined design rules, StreptoCAD will enable results to be fed back into the platform, allowing researchers to learn with each design iteration.

The next major bottleneck in *Streptomyces* genetic engineering is automating the practical laboratory work required to implement the designs generated by tools like StreptoCAD. Addressing this challenge would further streamline the workflow from design to execution, significantly accelerating research and development in this field. We are actively working on integrating laboratory automation protocols with StreptoCAD to overcome this hurdle. In its current implementation, the developed CAD tool is tuned for use in *Streptomyces*, however, with small modification it can easily be extended to be used beyond *Streptomyces*. This flexibility positions StreptoCAD as a versatile platform potentially benefiting a broader spectrum of microbial genetic engineering applications.

## Materials and Methods

### Web user interface

The StreptoCAD web interface was developed with the fully Python-based Plotly Dash^54^ and packaged as a Docker image (<u>deploy_apprunner_container.yml</u>). Hosted on AWS App Runner, the application ensures reliable and scalable access for researchers worldwide. The service can be found at https://streptocad.bioengineering.dtu.dk/.

### Code and data availability

StreptoCAD is an open-source project released under the MIT license. The source code is available on GitHub at https://github.com/hiyama341/streptocad. All data related to this study, including experimental results, protocols, and supplementary materials, can be accessed at https://github.com/hiyama341/streptocad/tree/main/notebooks/wet_lab_notebooks. These repositories provide comprehensive resources for reproducing the experiments and further exploring the capabilities of StreptoCAD.

### Modules of StreptoCAD

The core capabilities are driven by a set of Python functions and classes that are aimed at generating output from cloning (**<u>cloning module</u>**), generating and filtering sgRNAs (**<u>crispr</u> <u>module</u>**), generating and analyzing primers (**<u>primers module</u>**), loading in sequences (**<u>sequence_loading module</u>**), and finally, a module focused at wet lab functionalities (**<u>wet_lab module</u>**)(**<u>01-Construction_of_pOEX_PkasO_capture_plasmid.ipynb</u> & <u>02-Integration_of_GÖ4010_regulators_into_pOEX_PkasO.ipynb</u>)**. Utility functions are found in the root level folder with functions for easy data conversion between formats. All functions and classes are accessible with Python (>=3.9) or Jupyter Notebooks for ease of use. The documentation for how all functions work is available at ReadTheDocs at https://streptocad.readthedocs.io/en/latest/.

### sgRNA detection and filtering

The first step in sgRNA detection and filtering within StreptoCAD involves identifying potential sgRNA candidates across specified genomic regions or locus tags. The script <u>guideRNAcas3_9_12.py</u> first scans genomic sequences for PAM motifs corresponding to the selected CRISPR-Cas system (Cas3: ”**TTC**”, or Cas9: “**NGG**”). Potential sgRNAs are identified by their proximity to these motifs, and their sequences are then analyzed for off-target hits using a specified seed region (default: 13 bp). The sgRNAs are further filtered based on user-defined criteria, including GC content thresholds, specific PAM sequences, number of allowed off-targets, and downstream sequence motifs. The filtered sgRNAs are compiled into a DataFrame, and sorted by off-target count.

### Cytosine base-editing

The StreptoCAD pipeline also supports base-editing by identifying sgRNAs with editable cytosines. The code uses the <u>guideRNAcas3_9_12.py</u> script to find sgRNA sequences and then searches f for C-to-T conversion sites, annotating these potential base-editing sites with the <u>cripsr_best.py</u> script. The identified sgRNAs are then filtered to retain only those suitable for base-editing. In the final step, the base-editing sites are processed to identify and translate the resulting amino acid changes. As an extra feature if the switch “Editing Sequence Context” is switched on sgRNAs with an immediate guanine upstream of an editable cytosine are filtered out (https://github.com/hiyama341/streptocad/blob/main/streptocad/crispr/crispr_best.py).

### CRISPR interference

The StreptoCAD pipeline includes functionality for CRISPRi, allowing the identification and filtering of sgRNAs suitable for gene repression. The code uses the <u>guideRNA_crispri.py</u> to find sgRNAs targeting promoter regions and annotates their positions relative to the transcription start site (TSS). After identifying potential sgRNAs, the sequences are filtered based on criteria such as strand orientation, GC content, and proximity to the TSS (https://github.com/hiyama341/streptocad/blob/main/streptocad/crispr/guideRNA_crispri.py).

### *In silico* plasmid assembly and simulation

All plasmid assemblies within StreptoCAD are performed by simulating molecular biology techniques using key functions and classes from open-source libraries, including Pydna^55^, Biopython^56^, teemi^57^, Primer3^58^, and Pandas^59^. PyDNA and teemi facilitate DNA sequence manipulation and simulation, while Biopython provides tools for biological computation, Primer3 provides automated oligo analysis, and Pandas is employed for data manipulation and analysis. The code leverages these tools to design and simulate the construction of plasmids *in silico*, mimicking processes such as restriction enzyme digestion, ligation, and PCR amplification. Teemi is utilized for calculating primer annealing temperatures, which utilizes the NEB tm calculator^60^, optimizing PCR mixes, and managing sequence data. This approach ensures that all constructs are computationally validated before experimental work, streamlining the cloning process and optimizing workflows for complex genetic engineering tasks.

### Validation primers

Primers for validating the intended genome engineering were generated using our <u>primer_generation.py</u> script that automates the primer design process. For each specified locus tag or region, the script identifies the corresponding sequence from the GenBank file and extracts a target sequence, which includes a specified flanking region (default = 500 bp) around the provided sequence. Primers were designed using this target sequence to achieve a specified melting temperature (Tm). The script iteratively adjusts the flanking region size (default = 50 bp) if the designed primers do not yield a unique PCR product or exhibit undesirable characteristics such as significant hairpin formation or self-dimerization. This process continues until suitable primers are identified or a maximum number of iterations is reached (max_iter = 100).

### Requirements

Using the web app version of StreptoCAD requires only an internet connection. For users without internet access, the software can be deployed locally on a computer or run through Jupyter notebooks https://github.com/hiyama341/streptocad/tree/main/notebooks/app_workflows. Detailed package requirements for local deployment as a Docker image or through a virtual environment can be found here: https://github.com/hiyama341/streptocad/blob/main/README.md.

### Strains and Culture Conditions

All cloning procedures utilized One Shot Mach1 T1 Phage-Resistant Chemically Competent *E. coli* cells (ThermoFisher Scientific Inc., USA)^61^. These cells were grown on LB agar plates (10 g/L tryptone, 5 g/L yeast extract, 5 g/L sodium chloride, 15 g/L agar, brought to 1 L with MilliQ water) supplemented with either 50 µg/μL apramycin or 25 µg/μL chloramphenicol, and incubated at 37°C. For liquid cultures, *E. coli* was cultivated in 2xYT medium (16 g/L tryptone, 10 g/L yeast extract, 5 g/L sodium chloride, brought to 1 L with MilliQ water) in 15 mL culture tubes at 37°C with shaking at 250 rpm.

The constructed *Streptomyces* strains were cultured on mannitol soy flour (MS) agar plates (20 g/L fat-reduced soy flour, 20 g/L mannitol, 20 g/L agar, brought to 1 L with tap water) with 10 mM MgCl_2_ for conjugation experiments and spore production. Exconjugants were transferred to ISP2 agar plates (4 g/L yeast extract, 10 g/L malt extract, 4 g/L dextrose, 20 g/L agar, 333 mL tap water, 667 mL MilliQ water) supplemented with 50 µg/μL apramycin and 12.5 µg/μL nalidixic acid, and incubated at 30°C.

For liquid cultivation of *Streptomyces*, 350 mL shake flasks containing 50 mL of ISP2 medium, supplemented with 12.5 µg/μL apramycin, were used. Cultures were maintained in shaking incubators at 30°C and 180 rpm.

Drop-tests were performed on two types of media to evaluate phenotypic variations: DNPM media (40 g/L Dextrin, 7.5 g/L Peptone, 5 g/L Yeast Extract, 21 g/L MOPS, pH adjusted to 8 and brought to 1 L with MilliQ water) supplemented with 12.5 µg/mL apramycin, and MS media supplemented with 12.5 µg/mL apramycin. *Streptomyces* colonies were inoculated on these plates and incubated at 30°C.

### Genomic DNA Extraction, DNA concentration measurements, and Sanger sequencing

Genomic DNA extractions were carried out using DNeasy PowerSoil Pro Kits (Cat No./ID: 47014, Qiagen). The quality of the extracted DNA was assessed using a NanoDrop 2000 spectrophotometer (Thermo Fisher Scientific Inc., USA) to ensure high purity and concentration suitable for downstream applications.

Plasmid verification was performed by Sanger sequencing. For this purpose, plasmids were prepared and submitted using Mix2Seq overnight kits (Eurofins Genomics Germany GmbH, Germany). The sequencing data was analyzed using Benchling’s Sequence Alignment tool^62^ with StreptoCAD’s autogenerated plasmid sequences.

### Conjugations

The method for plasmid conjugation into *Streptomyces* was adapted from Whitford *et al.*, 2023^16^. In this approach, chemically competent *E. coli* ET12567 pUZ8002 cells were transformed with the target plasmid. The transformed cells were then spread on LB plates containing apramycin (50 µg/μL), chloramphenicol (25 µg/μL), and kanamycin (50 µg/μL) to select for transformants. After incubation at 37°C, the colonies were scraped off the plate using L-spreaders and resuspended in LB medium for overnight cultures.

Following overnight growth, the cultures were washed twice with LB medium to remove residual antibiotics and mixed with 500 μL spore suspension of the target *Streptomyces* strain. This mixture was then plated on MS (20 g/L D-Mannitol, 20 g/L Soy flour, 20 g/L Agar, 10 mM MgCl_2_) plates. After one day of cultivation (incubator at 30°C), the plates were overlaid with 1 mL of ddH_2_O and 5 μL of a 50 mg/mL apramycin solution, and incubated for an additional 4-5 days under the same conditions. Exconjugants were subsequently selected and transferred to ISP2 plates supplemented with 50 μg/ml apramycin for further culturing at 30°C. Spore preparation for the individual *Streptomyces* strain was carried out according to the procedure described by Tong *et al.*^15^.

### Plasmid Construction of pOEX-PkasO* platform

All primers used for plasmid construction were ordered from Integrated DNA Technologies (IDT). Primers were initially resuspended to a concentration of 100 µM using MilliQ water and subsequently diluted to a working concentration of 10 µM.

To develop the pOEX-PkasO* platform to allow for efficient cloning of regulators into an integrative vector, the existing pRMe4 plasmid was modified to include a P*KasO*\* and RBS region (“**GGAGG**”) flanked by a *StuI* restriction site for easy linearization in Hifi DNA Assemblies (Vector map: **S10**; Vector sequence **S11**.). Detailed protocols for the PCR reactions, gel purification, and assembly procedures are available in the supplementary **<u>01-Construction_of_pOEX_PkasO_capture_plasmid.ipynb</u>**. In short, the Hifi Assembly mixtures were incubated at 50°C for 1 hour. Subsequently, 5 µL of the assembly mixture was transformed into One Shot™ Mach1™ T1 Phage-Resistant Chemically Competent *E. coli*^61^ cells via heat-shock following the manufacturer’s protocol^61^, and the transformants were plated on LB + apramycin plates. Colonies were screened via diagnostic colony PCR (**<u>01-Construction_of_pOEX_PkasO_capture_plasmid.ipynb</u>**). DNA from positive colonies were then purified using the NucleoSpin Plasmid EasyPure kit^63^ (Machery-Nagel, Item number: 740727.50). The plasmid constructs were verified through Sanger sequencing using the Mix2Seq overnight kits (Eurofins Genomics, Germany) (**<u>01-Construction_of_pOEX_PkasO_capture_plasmid.ipynb</u>**).

### Plasmid Construction of Regulator Library

To develop the overexpression library, we PCR-amplified the coding sequence of the 18 selected regulators from the genomic DNA (gDNA) of *Streptomyces* Gö40/10, incorporating tails compatible with HiFi assembly. The plasmid backbone was linearized using *StuI* (**<u>02-Integration_of_GÖ4010_regulators_into_pOEX_PkasO.ipynb</u>**).

The HiFi assembly mix was then prepared and heat-shock transformed into the One Shot™ Mach1™ T1 Phage-Resistant Chemically Competent *E. coli* according to manufacturers protocol^61^ (**<u>02-Integration_of_GÖ4010_regulators_into_pOEX_PkasO.ipynb</u>**). Following transformation, we performed colony PCR to verify the correct transformants using our validation primers (Forward primer: 5‘-GCGGTGTTGTAAAGTCGTGGCC-3’, Reverse primer: 5’-CCGATCAACCGCGACTAGCATCG-3’). Positive candidates were selected and sequenced. Among the correctly assembled candidates, we grew overnight cultures and mini-prepped the plasmid DNA using the NucleoSpin Plasmid EasyPure kit^63^ (Machery-Nagel, Item number: 740727.50). This DNA was subsequently electroporated into *E. coli* strain ET122567, which facilitates the integration of the integrative plasmid into the *Streptomyces* Gö40/10 genome upon conjugation (**<u>02-Integration_of_GÖ4010_regulators_into_pOEX_PkasO.ipynb</u>**).

For the pre-culture, *E. coli* ET122567 was inoculated in 5 mL of YT medium containing 5 µL kanamycin (50 µM), 5 µL chloramphenicol (25 µM), and 5 µL culture from a cryotube, and incubated overnight in a shaker, at 250 rpm, 37°C. The conjugation was performed using the protocol detailed above. After conjugation, colonies were picked and transferred onto ISP2 plates supplemented with apramycin (**<u>02-Integration_of_GÖ4010_regulators_into_pOEX_PkasO.ipynb</u>**). Colony PCRs, using the validation primers (Forward primer: 5‘-GCGGTGTTGTAAAGTCGTGGCC-3’, Reverse primer: 5’-CCGATCAACCGCGACTAGCATCG-3’). were performed to verify successful integration. Confirmed colonies were streaked on ISP2 plates and spore suspensions were prepared as described above.

### Imaging and illustrations

All imaging was performed using a Nikon D7500 DSLR camera. The camera was controlled using the accompanying Nikon software, which allowed for precise adjustment of settings and consistent image capture. Figures were created using Adobe Illustrator.

## Abbreviations

CRISPR: clustered regularly interspaced short palindromic repeats
CBE: cytosine base editing
sgRNA: single guide RNA
BGCs: Biosynthetic gene clusters **CAD** - computer-aided design **DBTL** - Design-Build-Test-Learn
FAIR: Findable, Accessible, Interoperable, and Reusable
DSB: Double-strand break
UGI: Uracil glycosylase inhibitor
ssDNA: Single-stranded DNA
CRISPRi: CRISPR interference
TSS: Transcriptional start site
PAM: Protospacer adjacent motif
IDT: Integrated DNA Technologies

## Author Information

<u>Corresponding Authors</u> Rasmus J. N. Frandsen - Department of Biotechnology and Biomedicine, Technical University of Denmark, Kgs. Lyngby, Denmark; https://orcid.org/0000-0002-3799-6062;

**Tilmann Weber** - *The Novo Nordisk Foundation Center for Biosustainability, Technical University of Denmark, 2800 Kgs. Lyngby, Denmark*; https://orcid.org/0000-0002-8260-5120;

## Authors

**Lucas Levassor** - *The Novo Nordisk Foundation Center for Biosustainability, Technical University of Denmark, 2800 Kgs. Lyngby, Denmark;* Department of Biotechnology and Biomedicine, Technical University of Denmark, Kgs.

Lyngby, Denmark https://orcid.org/0000-0002-3261-9911

***Christopher M. Whitford*** *-* The Novo Nordisk Foundation Center for Biosustainability, Technical University of Denmark, 2800 Kgs. Lyngby,

## Denmark

https://orcid.org/0000-0001-5023-0830

**Søren D. Petersen** - Department of Biotechnology and Biomedicine, Technical University of Denmark, Kgs. Lyngby, Denmark https://orcid.org/0000-0003-4104-5144

**Kai Blin** - *The Novo Nordisk Foundation Center for Biosustainability, Technical University of Denmark, 2800 Kgs. Lyngby, Denmark* https://orcid.org/0000-0003-3764-6051

## Author Contributions

The project was conceptualized and designed by L.L., C.M.W., T.W, and R.J.N.F. Laboratory experiments were performed by L.L. and C.M.W., who also analyzed the data. The manuscript was drafted by L.L., C.M.W., S.D.P., R.J.N.F., and T.W., with contributions and feedback from all authors. Figures were prepared by C.M.W., with input from all authors. L.L. wrote the code with input from K.B.

## Acknowledgments

This research was funded by a grant from the Novo Nordisk Foundation (NNF20CC0035580).

## Notes

https://streptocad.bioengineering.dtu.dk/

### Competing Interest Statement

The authors have declared no competing interest.

