## Supplementary Material for "StreptoCAD: An open-source software toolbox automating genome engineering workflows in streptomycetes"

**Tables:**

- **Supplementary table 1.** All transcriptional regulators from LuxR and SARP families
- **Supplementary table 2.** Successfully integrated transcriptional regulators from LuxR and SARP families

**Figures:**

- **Supplementary figure 1.** Guide to using Workflow 1: Overexpression plasmid generation
- **Supplementary figure 2.** Guide to using Workflow 2: CRISPR-BEST
- **Supplementary figure 3.** Guide to using Workflow 3: Multiplexed CRISPR-BEST
- **Supplementary figure 4.** Guide to using Workflow 4: CRISPRi plasmid generation
- **Supplementary figure 5.** Guide to using Workflow 5: CRISPR-Cas9 plasmid generation
- **Supplementary figure 6.** Guide to using Workflow 6: CRISPR-Cas3 plasmid generation
- **Supplementary figure 7.** Plasmid map of pOEX-PkasO
- **Supplementary figure 8.** Diagnostic colony PCR of Streptomyces sp. Gö40/10 strains

**Supplementary files:**

- **Supplementary file 1**. Plasmid sequence of pOEX-PkasO* (FASTA)
- **Supplementary file 2**. antiSMASH 7 output for S. Gô40/10
- **Supplementary file 3.** Workflow 1-6 videos

**Supplementary table 1**. All transcriptional regulators from LuxR and SARP families

| locus_tag | region | regulator_# | description |
|---|---|---|---|
| LLPMBPKK_00292 | region 1 | Reg 1 | LuxR family DNA-binding response regulator |
| LLPMBPKK_00328 | region 1 | Reg 2 | LuxR family transcriptional regulator |
| LLPMBPKK_00586 | region 4 | Reg 3 | LuxR family DNA-binding response regulator |
| LLPMBPKK_01488 | region 10 | Reg 4 | transcriptional regulator, SARP family |
| LLPMBPKK_02197 | region 12 | Reg 5 | transcriptional regulator, SARP family |
| LLPMBPKK_02209 | region 12 | Reg 6 | LuxR family DNA-binding response regulator |
| LLPMBPKK_02563 | region 13 | Reg 7 | LuxR family DNA-binding response regulator |
| LLPMBPKK_02633 | region 14 | Reg 8 | transcriptional regulator, SARP family |
| LLPMBPKK_02635 | region 14 | Reg 9 | LuxR family DNA-binding response regulator |
| LLPMBPKK_02637 | region 14 | Reg 10 | LuxR family DNA-binding response regulator |
| LLPMBPKK_02662 | region 14 | Reg 11 | transcriptional regulator, SARP family |
| LLPMBPKK_05992 | region 22 | Reg 12 | LuxR family DNA-binding response regulator |
| LLPMBPKK_06337 | region 27 | Reg 13 | transcriptional regulator, SARP family |
| LLPMBPKK_06422 | region 28 | Reg 14 | transcriptional regulator, SARP family |
| LLPMBPKK_06907 | region 30 | Reg 15 | LuxR family transcriptional regulator |
| LLPMBPKK_07531 | region 33 | Reg 16 | LuxR family transcriptional regulator |
| LLPMBPKK_07744 | region 36 | Reg 17 | LuxR family transcriptional regulator |
| LLPMBPKK_07949 | region 38 | Reg 18 | transcriptional regulator, SARP family |

**Supplementary table 2**. Successfully integrated transcriptional regulators from LuxR and SARP families

| locus_tag | region | regulator_# | description |
|---|---|---|---|
| LLPMBPKK_00292 | region 1 | Reg 1 | LuxR family DNA-binding response regulator |
| LLPMBPKK_00328 | region 1 | Reg 2 | LuxR family transcriptional regulator |
| LLPMBPKK_00586 | region 4 | Reg 3 | LuxR family DNA-binding response regulator |
| LLPMBPKK_01488 | region 10 | Reg 4 | transcriptional regulator, SARP family |
| LLPMBPKK_02197 | region 12 | Reg 5 | transcriptional regulator, SARP family |
| LLPMBPKK_02563 | region 13 | Reg 7 | LuxR family DNA-binding response regulator |
| LLPMBPKK_02635 | region 14 | Reg 9 | LuxR family DNA-binding response regulator |

| | | | |
|---|---|---|---|
| LLPMBPKK_02662 | region 14 | Reg 11 | transcriptional regulator, SARP family |
| LLPMBPKK_05992 | region 22 | Reg 12 | LuxR family DNA-binding response regulator |
| LLPMBPKK_06422 | region 28 | Reg 14 | transcriptional regulator, SARP family |
| LLPMBPKK_06907 | region 30 | Reg 15 | LuxR family transcriptional regulator |
| LLPMBPKK_07531 | region 33 | Reg 16 | LuxR family transcriptional regulator |
| LLPMBPKK_07949 | region 38 | Reg 18 | transcriptional regulator, SARP family |

**Supplementary figure 1**. Guide to using workflow 1: Overexpression plasmid generation

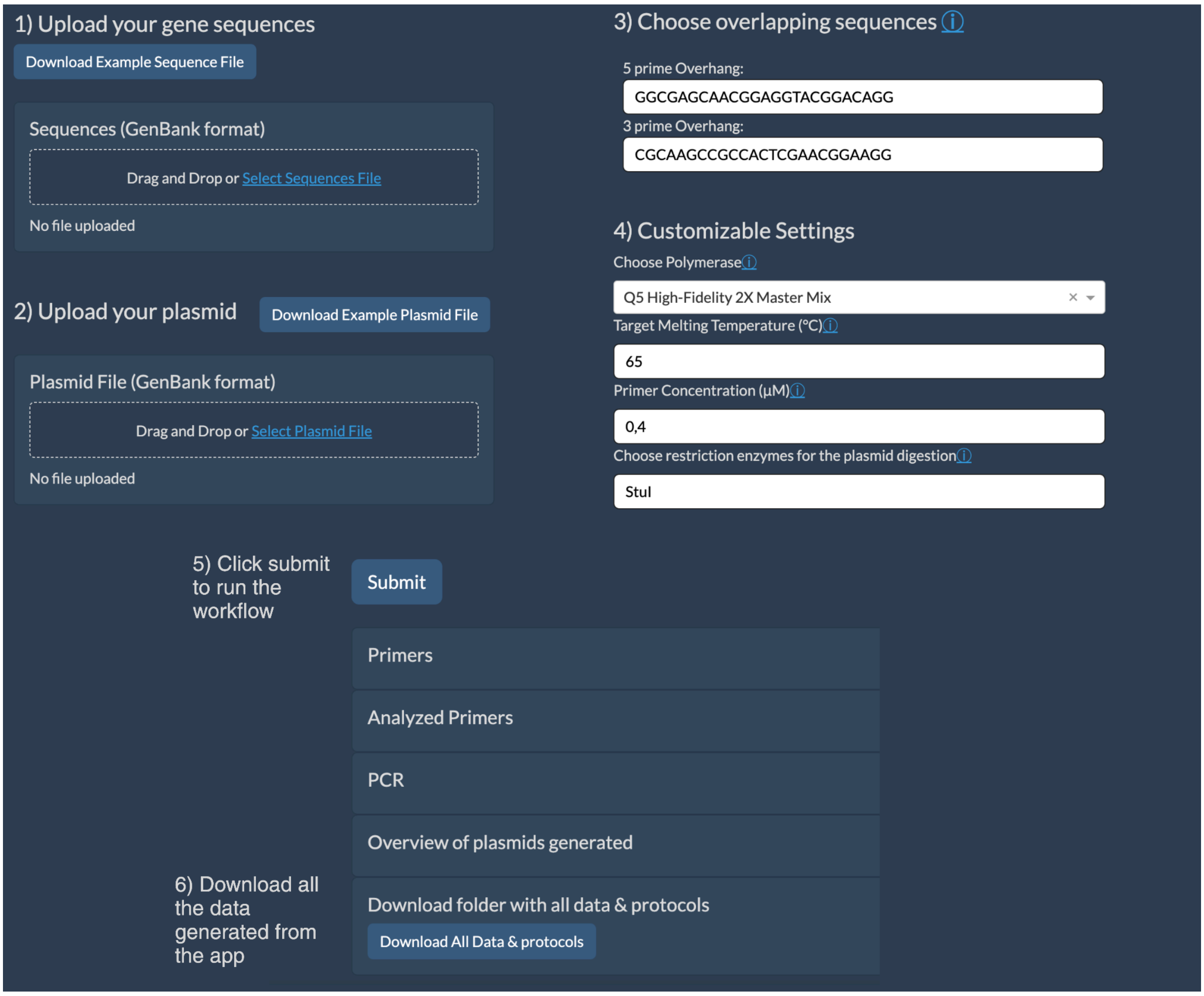


**Supplementary figure 1**: **Construction of an overexpression library. (1)** the user first uploads a GenBank file containing the desired targets for overexpression (example file: “Download example Sequence File”). **(2)** Next, the user uploads the pOEX-PkasO plasmid, available through "Download Example Plasmid File". **(3)** Users can then select overlapping sequences for up-and-down homology, with the default option configured to match pOEX-PkasO. **(4)** The user can also choose polymerase type, desired melting temperature, primer concentration, and starting number for the generated primers. **(5)** Clicking "Submit" starts the assembly process. **(6)** Once completed, the StreptoCAD platform displays the results, and the user can download all the results in by clicking the "Download all data & protocols" button which provides a zip file containing the input, and output data.

**Supplementary figure 2**. Guide to using Workflow 2 : CRISPR-BEST

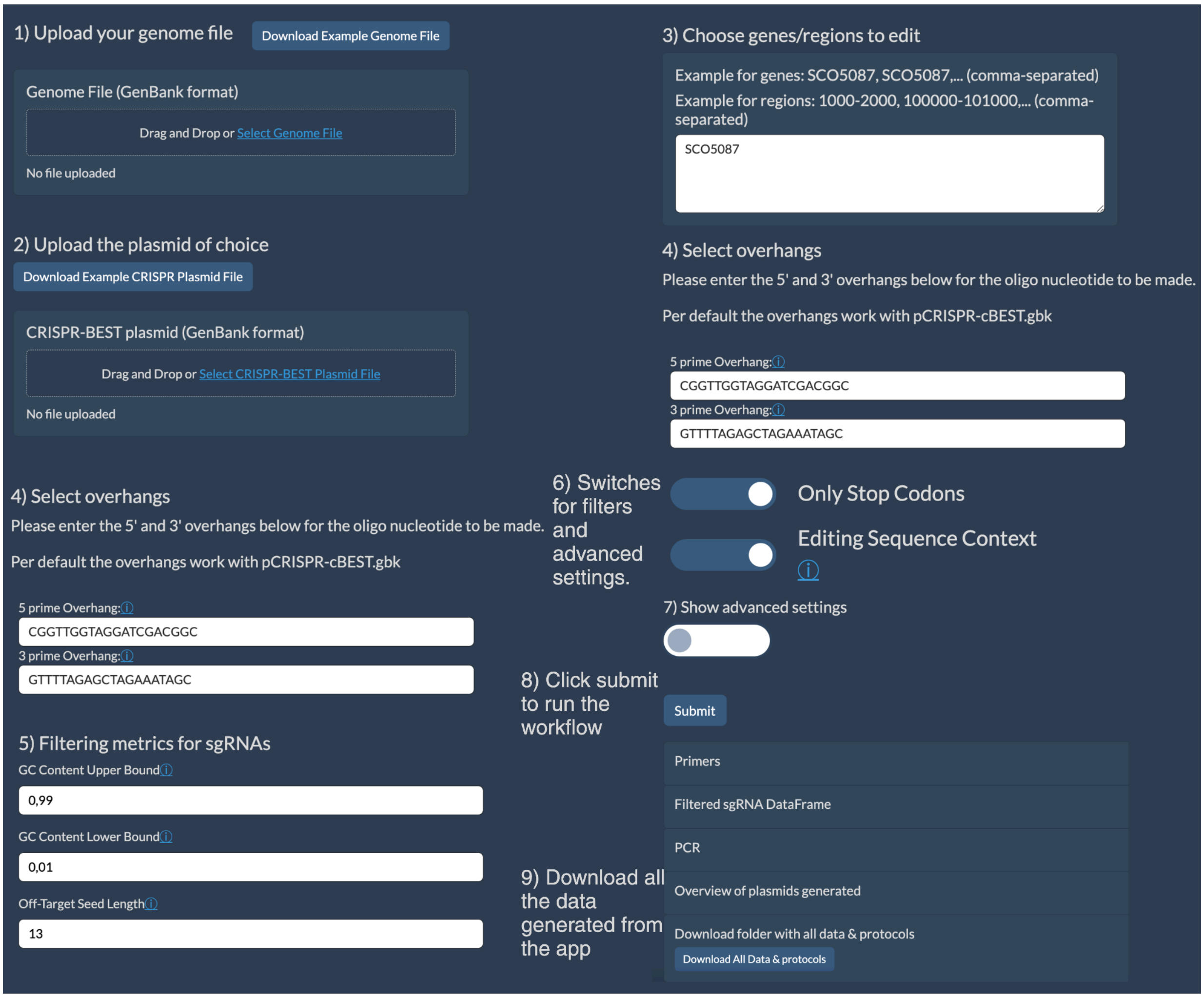


**Supplementary Figure 2**: **pCRISPR-cBEST workflow. (1)** The user begins by uploading the genome file, available through "Download example genome file" (S. coelicolor A3). **(2)** Next, the user uploads the pCRISPR-cBEST plasmid file, available through "Download example plasmid file (pCRISPR-cBEST.gb)". **(3)** The user specifies the genes to be base-edited by selecting from available locus tags, such as ‘SCO5087.’ **(4)** Users select overlapping sequences for up-and-down homology, with default settings matching the pCRISPR-cBEST plasmid along with sgRNA filtering metrics **(5)** An optional "Checking primers" checkbox can be ticked to automatically generate checking primers, and the user can also select the "Only Stop Codons" and "Sequence Editing Context" switches to refine the editing criteria. **(6)** Advanced polymerase settings make it possible to choose the polymerase, desired melting temperature, primer concentration, and validation primer distance from edit (default: 500 bp). **(7)** the user clicks "Submit" to initiate assembly. **(8)** When the computation is done the generated primers and plasmids are shown and, the "Download All Data & Protocols" button provides a zip file containing all relevant input and output data.

**Supplementary figure 3**. Guide to using Workflow 3 : Multiplexed CRISPR-BEST

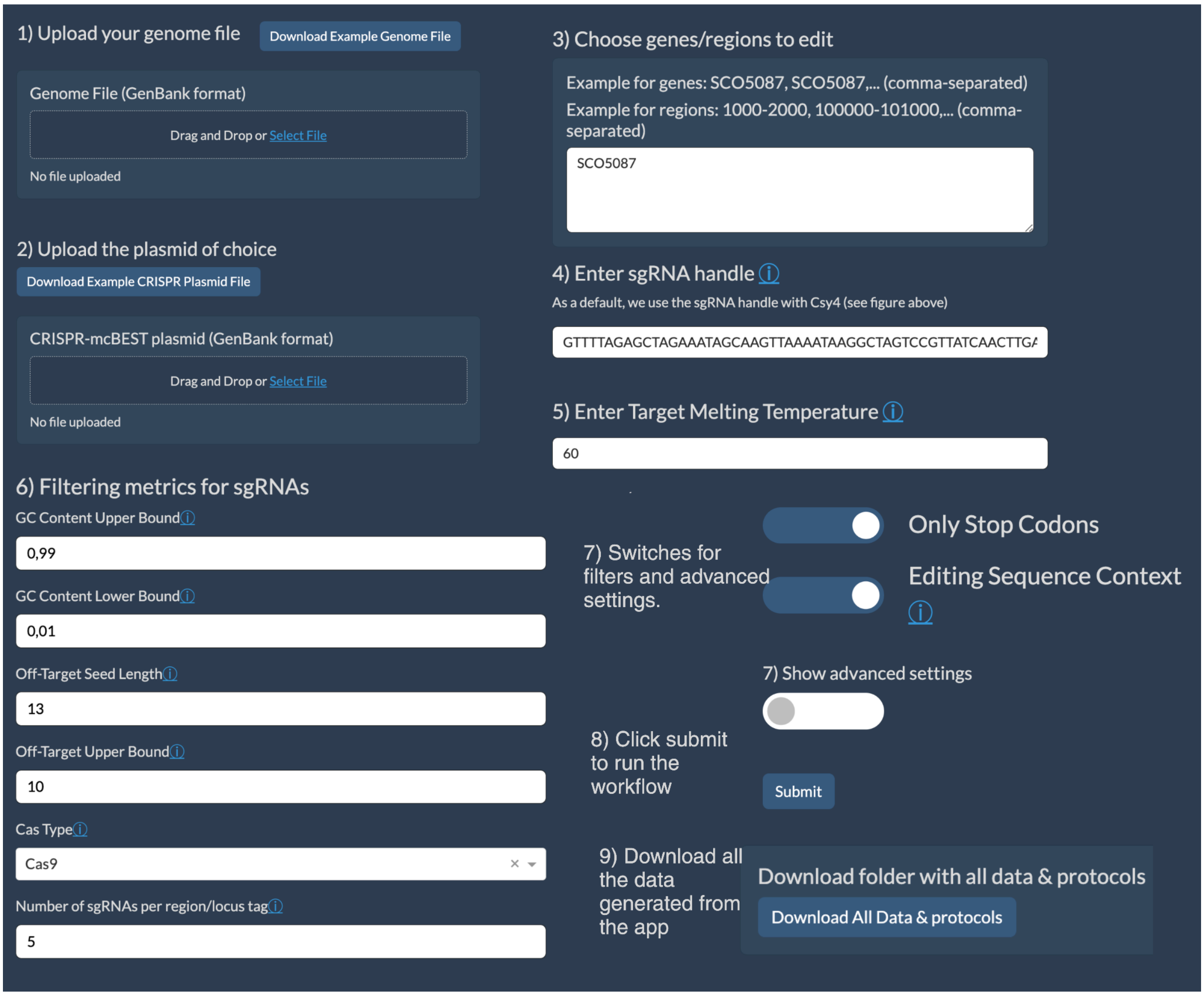


**Supplementary Figure 3: Multiplexed CRISPR-BEST sgRNA Integration Workflow. (1)** The user starts by uploading the genome file in GenBank format, accessible through "Download example genome file (S. coelicolor A3)". **(2)** Next, the user selects a plasmid file, either from their local system or by clicking on the provided example file link (pCRISPR-mcBEST.gb). **(3)** The user specifies the genes to be knocked out by selecting from available locus tags, such as SCO5087, SCO5089, and SCO5090. **(4)** Filtering metrics for sgRNAs, including off-target analysis and annealing sequences, are then configured according to the target genes. **(5)** The user chooses the polymerase type and target melting temperature to ensure optimal amplification. **(6)** After finalizing these settings, clicking "Submit" prompts StreptoCAD to design and arrange the sgRNAs into a multiplexed plasmid for CRISPR-BEST genome editing. **(7)** Upon completion, the platform displays the outputs, including primers and PCR schemes. **(8)** The "Download workflow" button provides a zip file with all relevant input and output data, supporting reproducibility and thorough documentation.

### **Supplementary Figure 4.** Guide to using Workflow 4: CRISPRi plasmid generation

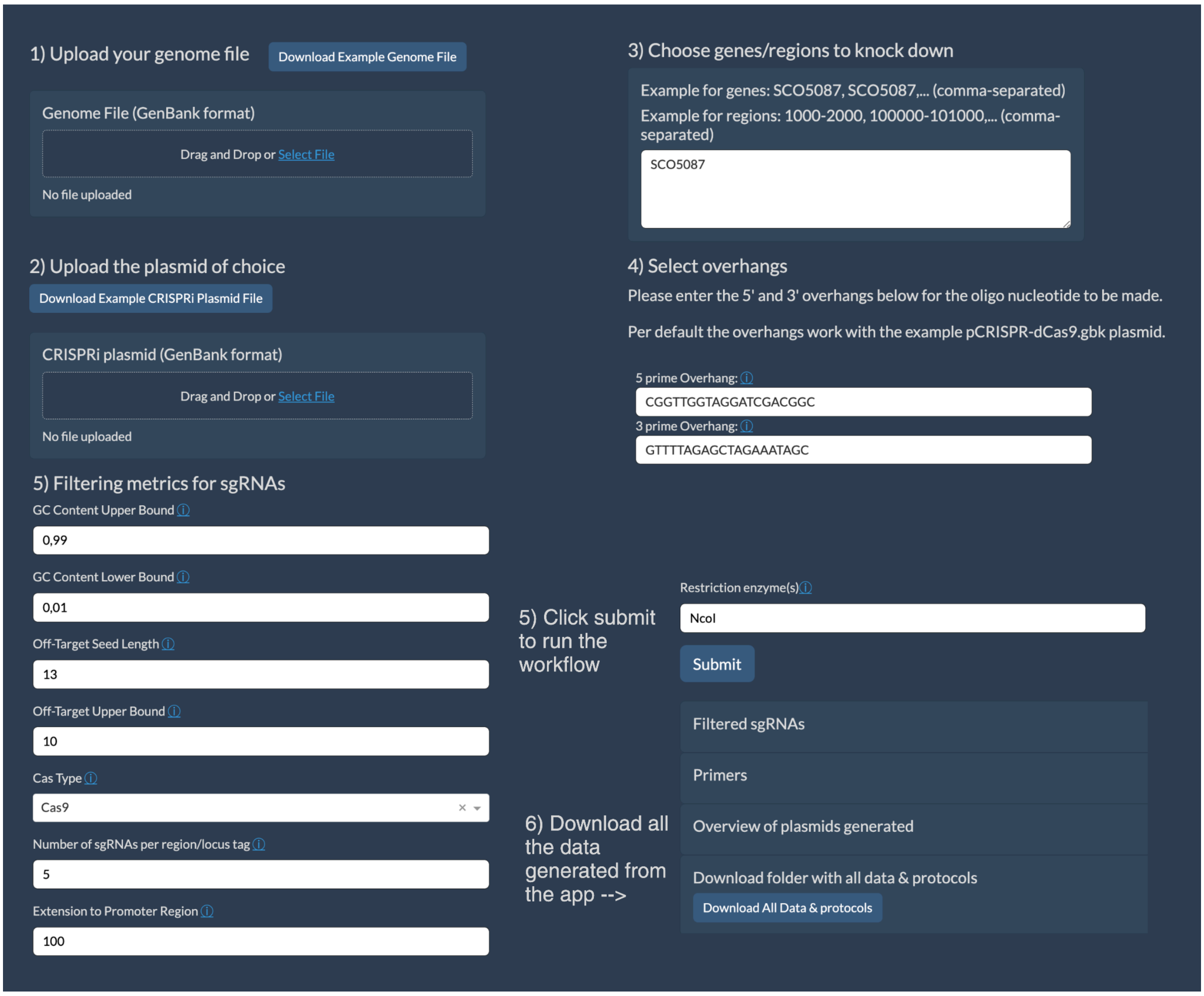


**Supplementary Figure 4: CRISPRi plasmid generation. (1)** The user begins by uploading a genome file in GenBank format, with example files available through "Download example genome file" (S. coelicolor A3). **(2)** Next, a relevant plasmid containing dCas9 and a cloning site is provided, either by uploading a custom plasmid file or by downloading the example pCRISPR-dCas9 plasmid via the "Download Example Plasmid File" button. **(3)** The user specifies the genes to be knocked down by selecting the appropriate locus tags, such as ['SCO5087', 'SCO5089', 'SCO5090'], and configures filtering metrics for sgRNAs and ssDNA overhang sequences to facilitate transcriptional interference. **(4)** The user then selects the polymerase type and sets the desired melting temperature. **(5)** Clicking "Submit" initiates the automated design process **(6)** The platform then displays the results, including a DataFrame with the sgRNAs, primers, and PCR schemes. **(7)** A "Download All Data & Protocols" button provides a zip file containing all input and output data.

**Supplementary Figure 5.** Guide to using Workflow 5: CRISPR-Cas9 plasmid generation

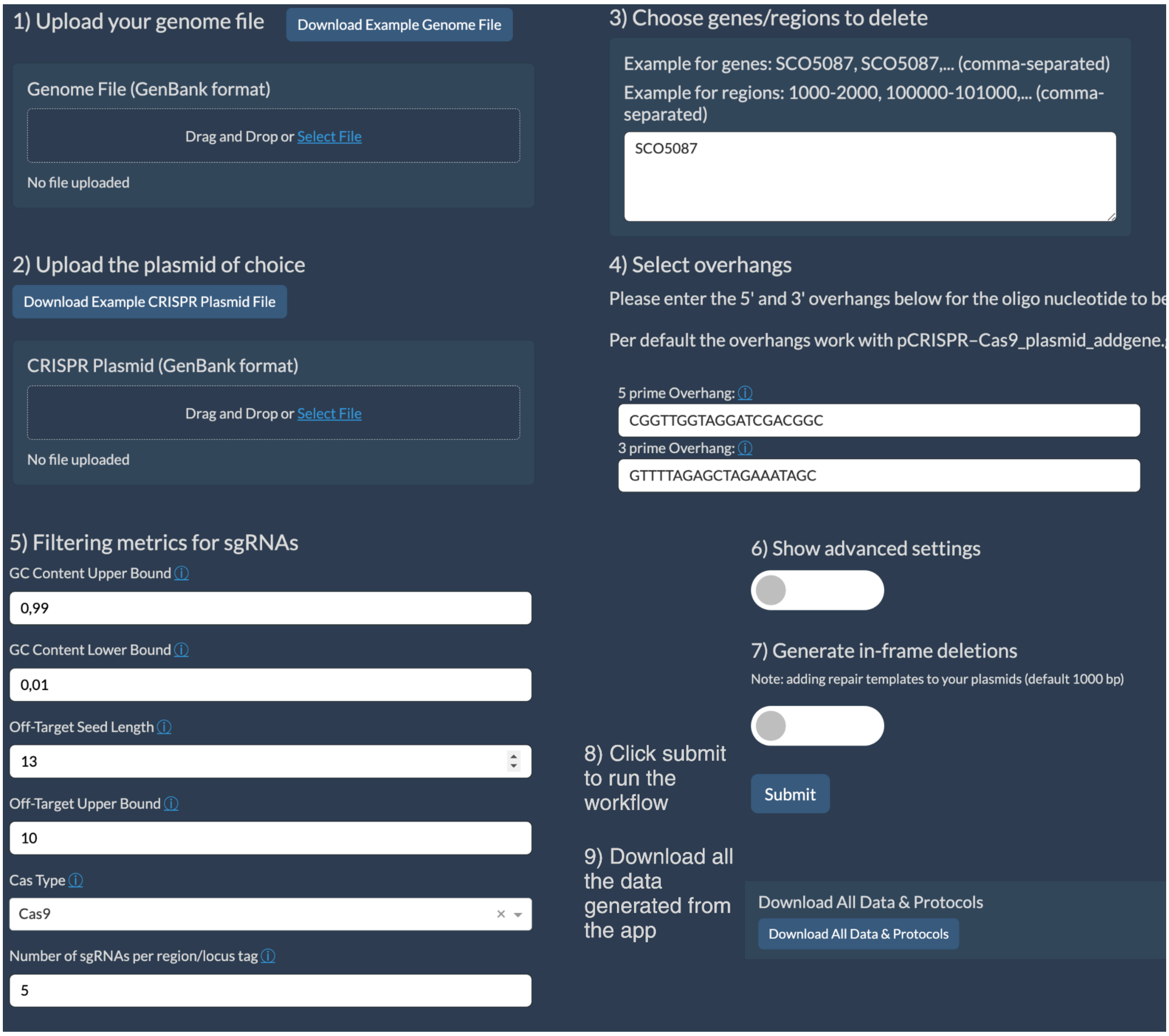


**Supplementary figure 5. Guide to using Workflow 5: CRISPR-Cas9 plasmid generation.** **(1)** The user begins by uploading the genome file of their choice in GenBank format. An example genome file for S. coelicolor can be accessed via "Download Example Genome File". **(2)** Next, the user selects a CRISPR-Cas9 plasmid file, which can be downloaded by clicking "Download Example Plasmid File”. **(3)** The user specifies the genes to knock out by choosing from available locus tags, such as locus tag: SCO5087, or by specifying a region, e.g., 10000-101000. **(4)** The user configures sgRNA filtering metrics, ssDNA overhang sequences (default settings match the pCRISPR-Cas9 plasmid), polymerase type, and desired melting temperature. **(5)** The "Generate In-Frame Deletion" switch can be activated to find repair templates up and downstream (default: 1000 bp) of the locus tag or region and to generate primers for Gibson cloning. **(6)** Clicking "Submit" initiates the design of sgRNAs and assembly of the plasmid, simulating ssDNA bridging, and if the "Generate In-Frame Deletion" switch is enabled, proceeds with Gibson cloning to integrate amplified upstream and downstream regions. **(7)** The platform displays the results, including primers for in-frame deletions, PCR schemes, and GenBank files. A "Download All Data & Protocols" button generates a zip file containing all input and output data, ensuring thorough documentation and reproducibility.

**Supplementary Figure 6.** Guide to using Workflow 6: CRISPR-Cas3 plasmid generation

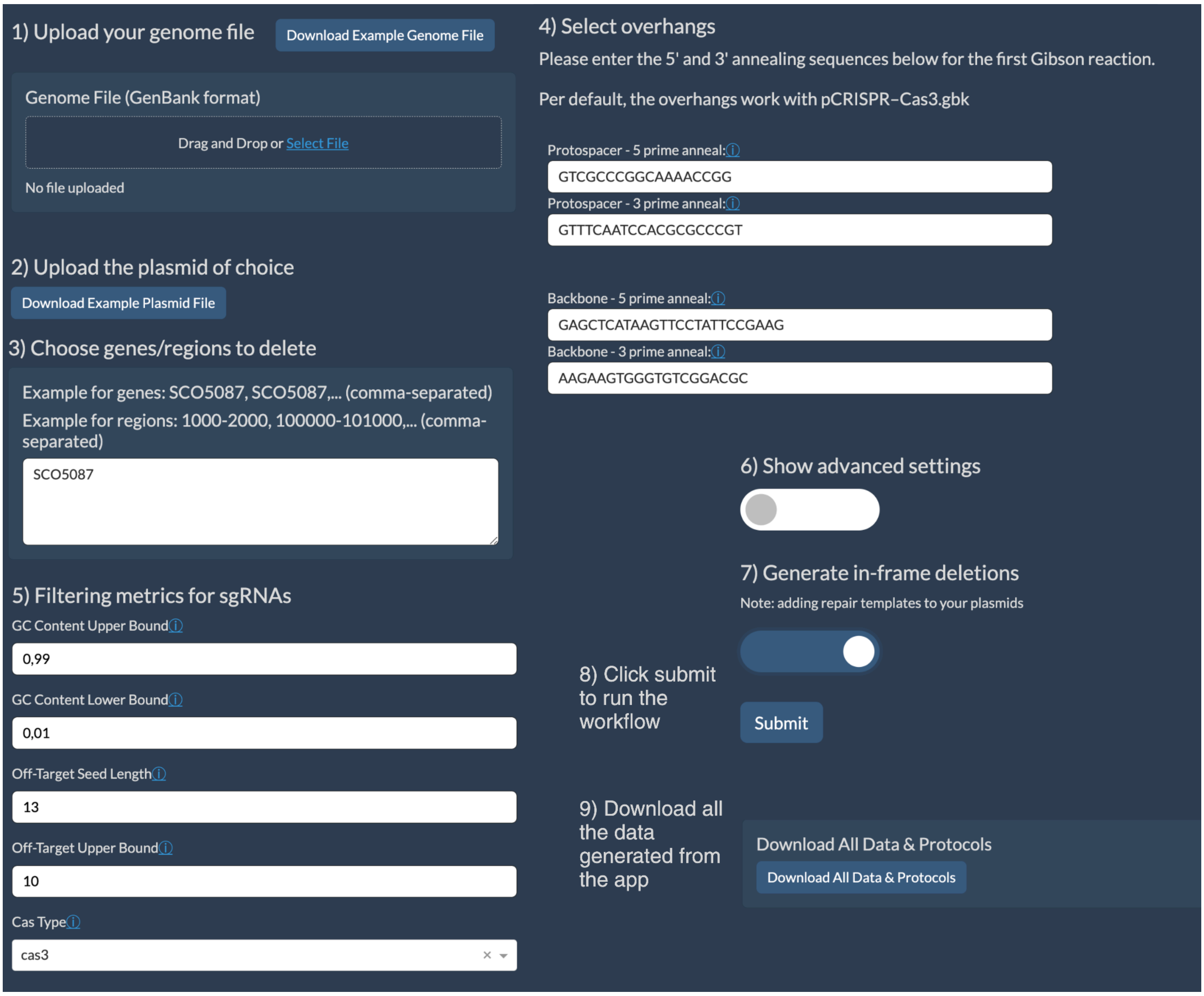


**Supplementary figure 6. Guide to using Workflow 6: CRISPR-Cas3 plasmid generation. (1)** The user begins by uploading the target genome file in GenBank format, with the option to access an example file for S. coelicolor through "Download Example Genome File". **(2)** Next, the user selects a CRISPR-Cas3 plasmid file (Click "Download Example Plasmid File".). **(3)** The user specifies the genes to delete by selecting the appropriate locus tags, such as ['SCO5087', 'SCO5089', 'SCO5090'], or regions (1000-10100, 90090-91090), and configures sgRNA filtering metrics. **(4)** ssDNA overhang sequences are selected (default matching the Cas3 system), as well as polymerase type and desired melting temperature. **(5)** The "Generate In-Frame Deletion" option can be activated to retrieve 1000 bp upstream and downstream sequences for in-frame deletion, and the software generates primers for these ssDNA oligos. **(6)** Clicking "Submit" initiates the design and simulation of the CRISPR-Cas3 workflow, including plasmid assembly using Gibson cloning. StreptoCAD then performs a secondary digestion with restriction enzymes (default: EcoRI), to complete the assembly. **(7)** The platform displays the results, including primers for in-frame deletions, and PCR schemes. **(8)** A "Download All

Data & Protocols" button generates a zip file containing all data needed for experimental use, including IDT order formats, repair templates, and assembly protocols, ensuring reproducibility and adherence to FAIR principles.

**Supplementary Figure 7:** Plasmid map of pOEX-PkasO*

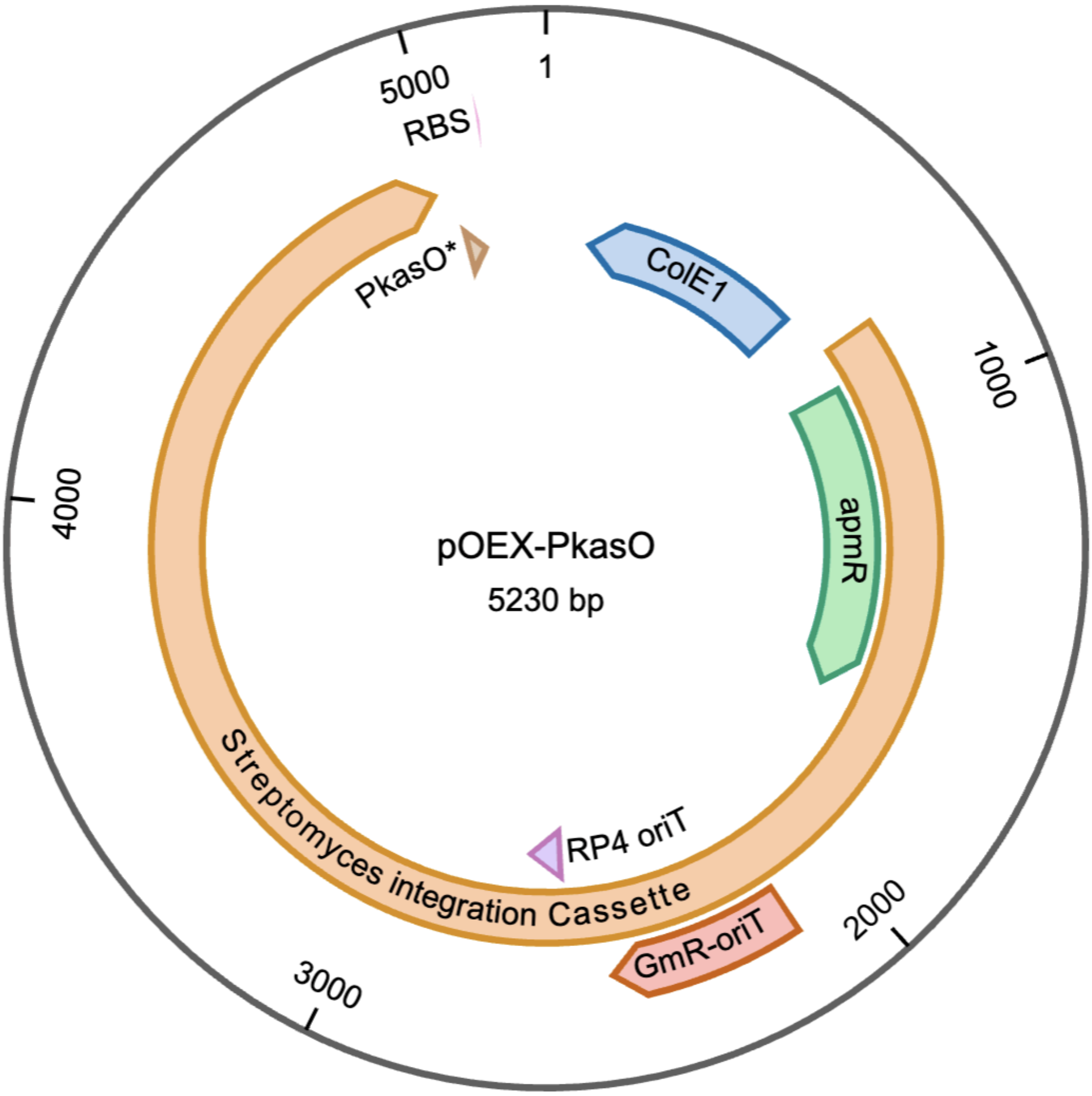


**Supplementary Figure 7: Plasmid Map of pOEX-PkasO.*** This plasmid, pOEX-PkasO*, is designed for the overexpression of target genes in Streptomyces. Built on the integrative pRM4 backbone, it includes a StuI restriction site for standardized gene insertion, allowing gene integration via PCR with defined overhangs and Gibson assembly. The plasmid features the strong Streptomyces promoter PkasO* and a canonical ribosome binding site (RBS/Shine Delgarno) sequence "GGAGG" to drive high-level transcription. Additionally, pOEX-PkasO* contains an apramycin resistance gene (apmR) for selection, as well as the PhiC31 integrase, enabling stable chromosomal integration at the attB site. The plasmid also harbors ColE1 and GmR origins of replication for maintenance in E. coli and

Streptomyces, and RP4 for conjugation.

### Supplementary Figure 8: Diagnostic colony PCR of *Streptomyces* sp. Gö40/10 strains

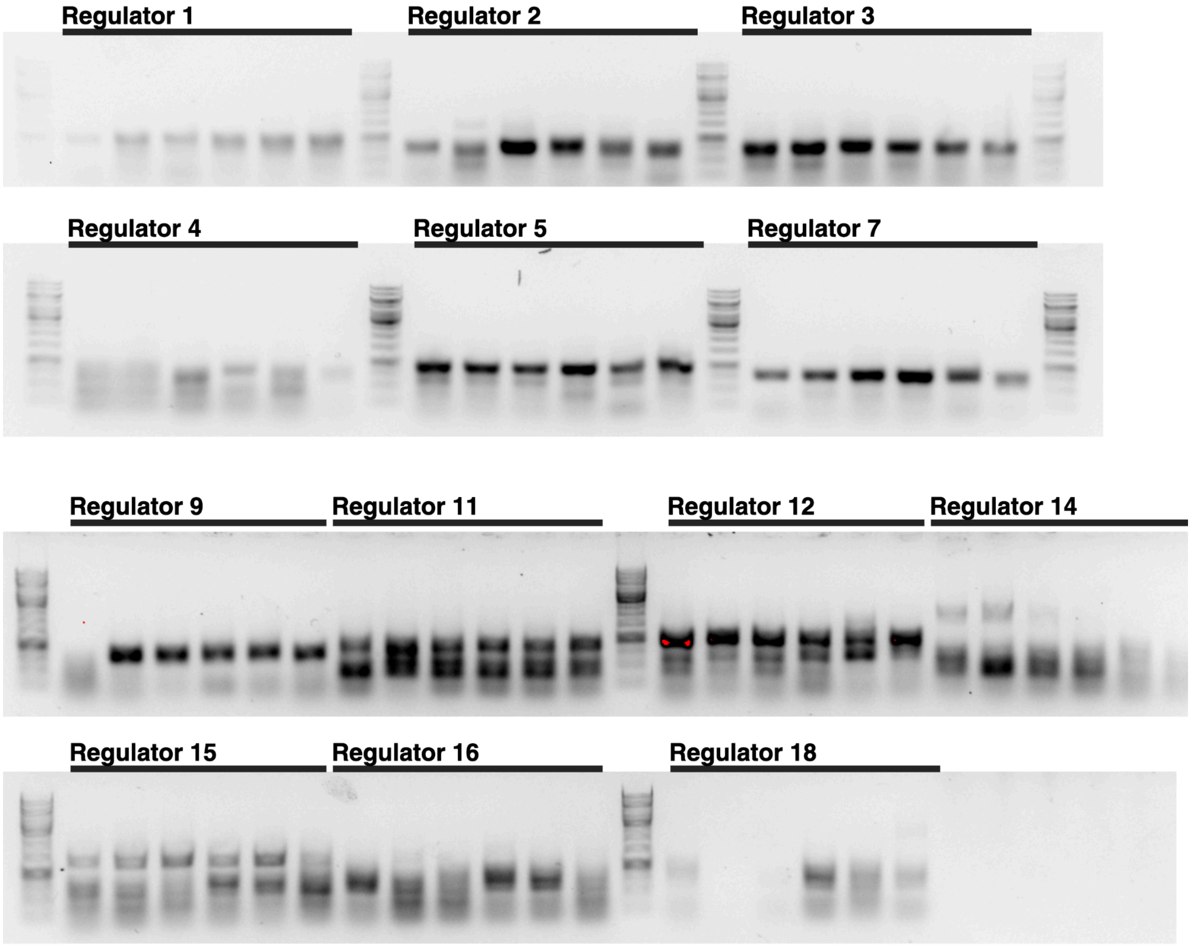


**Suppl. Figure 8. 1% Agarose gel of the Diagnostic PCR of the 13 regulators integrated into the *Streptomyces Gö40/10* genome.** Six colonies were picked from each strain, DNA was purified and PCRs were set up with our validation primers. 1kb ladder was used in between each regulators PCR products.

### **Supplementary File 1:** Plasmid sequence of pOEX-PkasO* (GenBank)

https://github.com/hiyama341/streptocad/blob/main/web_app/assets/pOEX-PkasO.gb.

**Supplementary File 2:** antiSMASH 7 output for *Streptomyces* sp. Gô40/10

antiSMASH 7 output for S. Gô40/10 download link

**Supplementary File 3:** Workflow 1-6 videos

https://github.com/hiyama341/streptocad/tree/main/workflow_videos
